# AdaReg: Data Adaptive Robust Estimation in Linear Regression with Application in GTEx Gene Expressions

**DOI:** 10.1101/869362

**Authors:** Meng Wang, Lihua Jiang, Michael P. Snyder

## Abstract

With the development of high-throughput RNA sequencing (RNA-seq) technology, the Genotype Tissue-Expression (GTEx) project (Consortium et al., 2015) generated a valuable resource of gene expression data from more than 11,000 samples. The large-scale data set is a powerful resource for understanding the human transcriptome. However, the technical variation, sequencing background noise and unknown factors make the statistical analysis challenging. To eliminate the possibility that outliers might affect the estimation of population distribution, we need a more robust estimation method, a method that will adapt to heterogeneous genes and further optimize the estimate for each gene. We followed the approach of the robust estimation based on *γ*-density-power-weight (Fujisawa and Eguchi, 2008; Windham, 1995), where *γ* is the exponent of density weight which controls the balance between bias and variance. As far as we know, our work is the first to propose a procedure to tune the parameter *γ* to balance the bias-variance trade-off under the mixture distributions. We constructed a robust likelihood criterion based on weighted densities in the mixture model of Gaussian population distribution mixed with unknown outlier distribution, and developed a data-adaptive *γ*-selection procedure embedded into the robust estimation. We provided a heuristic analysis on the selection criterion and found that our practical selection trend under various *γ*’s in average performance has similar capability to capture minimizer *γ* as the inestimable Mean Squared Error (MSE) trend from our simulation studies under a series of settings. Our data-adaptive robustifying procedure in the linear regression problem (AdaReg) shows a significant advantage in both simulation studies and real data application of heart samples from the GTEx project compared to the fixed *γ* procedure and other robust methods. This paper discusses some limitations of this method, and future work.

## 1 Introduction

The emergence of high-throughput RNA sequencing (RNA-seq) technology dramatically increases the development of gene expression analysis. The Genotype Tissue-Expression (GTEx) project (Consortium et al., 2015) generated the gene expression data from RNA-seq from more than 11, 000 samples of 53 tissues (up to the version 7 release from GTEx Portal), providing a valuable resource to study tissue variation in the human transcriptome. However, the technical variation, sequencing background noise and unknown factors make statistical analysis challenging. As an example, consider the expression for gene MYH7 (myosin heavy chain 7) in heart atrial appendage and heart left ventricle. Figure 1 (in the left panel) shows sample densities in these two parts of heart. We can see a long tail in the low expression end of the distribution of left ventricle samples. These outliers could be caused by technical noise, or could represent an abnormality in the sample expression. This is only for one gene, while different genes have different types of outliers. The outlier proportions can be large or small, and their magnitudes vary from gene to gene. Given the presence of various outliers, there is a great need to have a robust estimation method, especially adaptive to different cases.

**Figure 1:**
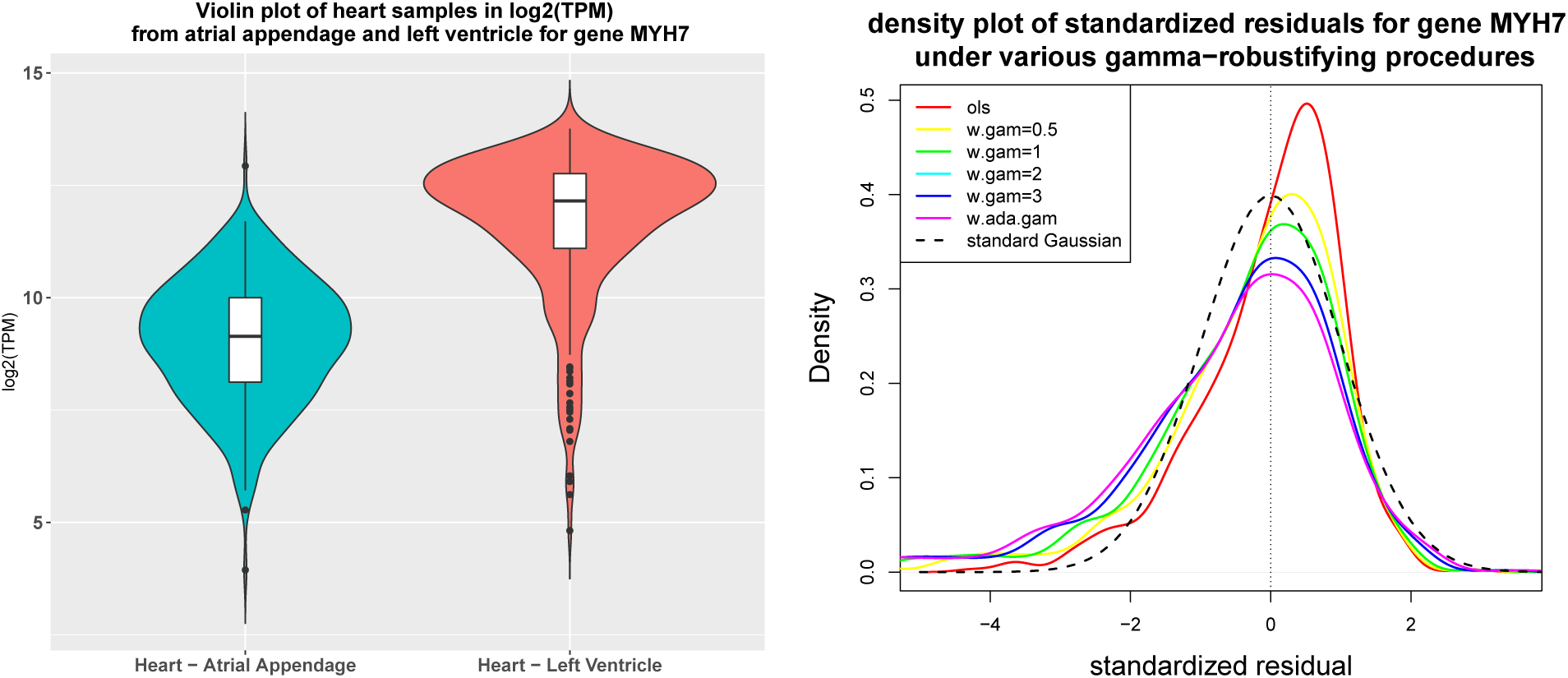
Motivation example of heart samples from atrial appendage and left vertical for gene MYH7 in GTEx RNA-seq data. The left panel is the violin plot of the heart expression in logTPM in two heart groups. The right panel is the density plot of standardized residuals (defined in (25)) from various *γ*-robustifying procedures and the dashed black curve is the standard Gaussian density without containing any outliers.

The literature of robust estimation in linear regression is very rich (Hampel et al., 2011; Huber, 2011; Maronna et al., 2018; Rousseeuw and Leroy, 1987). One approach is based on selecting a subset less influenced by the leverage points such as the least median squares (lms) (Rousseeuw, 1984), finding the narrowest hyperplanes covering half of the observations, and the least trimmed squares (lts) (Rousseeuw, 1984, 1985), removing the topmost most leverage points. The other approach is based on down-weighting the outliers by choosing various weight functions forming the M-estimate. Classical seminal works include Huber’s weight (Huber, 1964),, Hampel’s three-part weight, Tukey’s bisquare weight, S-estimation (Rousseeuw and Yohai, 1984), and others. Windham (1995) proposed a robustifying procedure based on the density power weight 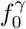 (*γ* ≥ 0) to robustly fit the population model *f*_0_ (extended details in Subsection 2.2), which is related to the density weight divergence in Basu et al. (1998). Jones et al. (2001) gave a comparison of these two methods. Fujisawa and Eguchi (2008) revisited these old works and constructed a *γ*-cross entropy robust criterion, assuming under a proper *γ*(≥ 0), the outliers go to the tails of density power 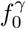 and thus do not contribute much in the population estimation. Recently, the *γ*-cross entropy criterion has gained much attention and there are a series of variant works including robust estimation using an unnormalized model (Kanamori and Fujisawa, 2015), robust clustering (Chen et al., 2014), Gaussian graphical modeling (Katayama et al., 2018; Miyamura and Kano, 2006), and others.

One useful property of both the *γ*-cross entropy criterion and the *γ*-density-power-weight is that the estimation procedure is controlled by only one system parameter: *γ*. Many previous papers pointed out that *γ* controls the trade-off between robustness and efficiency of the estimate. In Subsection 2.4, we investigate the trade-off in terms of bias and variance. However, how to choose a proper *γ* in practice is still unknown. Previous *γ*-robustifying procedures are based on a fixed preselected *γ*. In the example of gene MYH7, Figure 1 (in the right panel) shows the density plot of standardized residuals under various *γ*’s. When *γ* = 0, the estimation is the same as from Ordinary Least Squares (OLS). Ideally, if *γ* is the properly selected, the residual peak should be around zero. Under *γ* = 0, 0.5, 1, the residual peaks are obviously greater than zero, which means such *γ* is not large enough so that there is still some information left in the residuals, while under *γ* = 2, 3, the density peaks are close to zero. We can see this gene prefers a large *γ*. As we have pointed out, the outliers vary from gene to gene and thus a proper *γ* also varies among different genes. This motivates us to develop a procedure to select a proper *γ* to adapt to each gene.

Our contribution is that following the approach of using density power weight, we construct a robust weighted likelihood estimation criterion based on weighted densities in the mixture model, and develop a data-adaptive *γ*-selection procedure in linear regression (AdaReg). We provide a heuristic analysis on the selection criterion, and find that our practical selection trend under various *γ*’s in average performance has similar capability to capture minimizer *γ* as the inestimable Mean Squared Error (MSE) trend from our simulation studies under a series of settings. In this paper, we mainly discuss the estimation problem in linear regression. We believe our data-adaptive robustifying procedure will broaden the direction of density power weight in more applications.

The outline of the rest of the paper is as follows: In Section 2, we develop and analyze our algorithm, AdaReg. In Subsection 2.1, we first set up the regression problem in a mixture model. In Subsection 2.2, we consider weighting the model to purify the mixture. In Subsection 2.3, using our weighted mixture model, we construct our robust likelihood criterion based on the weighted densities, and provide robust estimation under a fixed *γ*. In Subsection 2.4, we investigate the bias and variance trade-off in terms of *γ*. We develop a novel *γ* selection criterion in Subsection 2.5. In Subsection 2.6, we summarize our data-adaptive algorithm (AdaReg) in the linear regression problem. Section 3 covers simulations and applications. We apply AdaReg in our simulation studies in Subsection 3.1, and in a real dataset of heart samples from RNA-seq in the GTEx project in Subsection 3.2. We compare our method to the fixed *γ* procedures and other robust regression methods. Finally, in Section 4, we summarize our work, point out a few limitations, and suggest some further work.

## 2 Method

### 2.1 Problem setup

Suppose we would like to investigate the relationship between a response variable **y** ∈ ℝ^*n*^ and design matrix **X** ∈ ℝ^*n×p*^ from *n* samples. For example, we are interested in investigating gene expression variation under different tissue types. In the similar notation as in (Katayama et al., 2018), consider the response variable **y** coming from a mixture model,

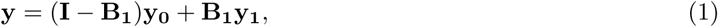

where **I** ∈ ℝ^*n×n*^ is the identity matrix and **B**_**1**_ is a diagonal matrix where the diagonal elements are Bernoulli variables with probability *π*_1_ ∈ [0, 0.5) to be 1, and **y**_**0**_ is the clean part forming the population, and **y**_**1**_ is the outlier. The 0-1 elements in the diagonal of **B**_**1**_ indicate the samples coming from the clean part **y**_**0**_ or the outlier **y_1_**. We assume independence among all the components of **y**. We only consider the outliers in the response variable not in the covariates here. Suppose the clean part of **y** comes from the ordinary linear model,

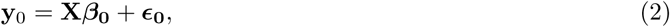

where ***β***_**0**_ ∈ ℝ^*p*^ is the coefficient vector, and the Gaussian independent noise 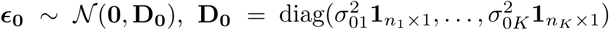, *K* is the group number, *n*_*k*_ is the sample size in group *k*, and each group can have its own variance 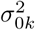. We further parameterize the population distribution as Gaussian. For the RNA-seq data, as a convention, we take logarithm transformation on the standardized expression of Transcripts Per Kilobase Million (TPM) or Reads Per Kilobase Million (RPKM) to make the density more symmetric, more Gaussian distributed. For the intensity data such as from microarray or mass spectrometry platform, the expression data is conventionally assumed as Gaussian distributed in the log scale. Under unknown distribution of the outliers and non-vanishing outlier proportion, our goal is to robustly estimate the population parameters 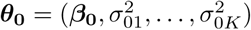 and *π*_0_ for each gene.

In the following analysis, we first consider there is only one group, i.e., the design matrix **X** is simply **1_n_**_*×***1**_, and develop a criterion to data-adaptively select *γ* then go back to embed our selection procedure into a general linear regression setting.

### 2.2 Weighted mixture density

Suppose the response **y** is from one group. Based on the setting (1)–(2), the problem is reduced to each component *y*_*i*_ from a mixture density

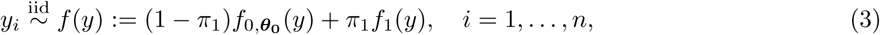

where *f* is the mixture density function, 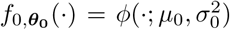 is for the population modeled as Gaussian density *φ* with mean *µ*_0_ and variance 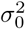, and *f*_1_ is the unknown outlier density.

In the early work, Windham (1995) considered to attach a weight 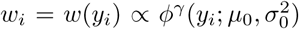 to each *y*_*i*_ where *γ* ≥ 0 is the exponent parameter and the weights are self-standardized, that is, 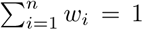. In this way, weighting the samples by the power of the population density, the points coming from the population gain more weights, while the outliers gain less weights so that the outliers do not contribute much in the population estimation. To estimate the score function *s*(*y, **θ***):= *∂* log *f*_0,***θ***_(*y*)*/∂**θ***, matching its empirical average from the weighted samples and the corresponding theoretical expectation gives the estimating equation for the Windham’s procedure,

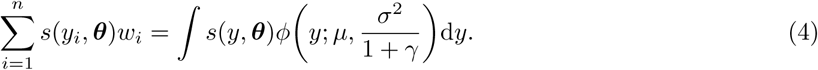

When *γ* = 0, the solution to (4) is the Maximum Likelihood Estimation (MLE).

Now we formulate the process of Windham’s procedure in terms of weighting the mixture density. Consider multiplying 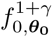 on both sides of (3), then re-standardizing each product to a density function gives our weighted mixture model,

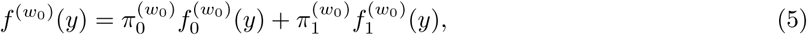

where

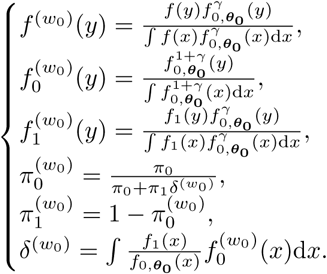

In the weighted mixture model (5), the function with superscript (*w*_0_) means the function depends on the weight 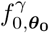. When the population and outlier distributions, the null and the alternative, are not hard to distinguish, under a proper *γ*, the integral of the likelihood ratio of *f*_1_ to 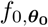 is small in the probability measure of the weighted null density. In such scenario, 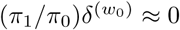 and thus 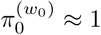. Therefore, our weighted mixture model can be approximately purified to

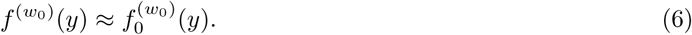

In the case that 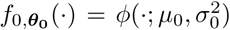, the empirical mass density of 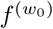 at *y*_*i*_ can be estimated by the weight

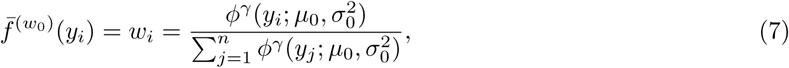

and 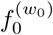 is the density of Gaussian distribution 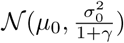 with mean *µ*_0_ and variance 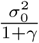, shrinking the variance to 1/(1 + *γ*) of the original variance, and thus the outliers go to the tails of the variance-shrunk density.

**Remark**: unlike other weighting procedures such as Huber, Hampel, Tukey’s methods where the weights are basically loss penalty, the weights in the *γ*-robustifiying procedure take the role of transforming samples from a mixture distribution to a more purified distribution.

### 2.3 Robust estimation under a fixed *γ*

From the estimation equation in (4), the Windham’s procedure essentially relies on the weighted density approximation (6). Here we take a direct approach, measuring the ordinary cross entropy of the weighted mixture density to the weighted theoretical null density,

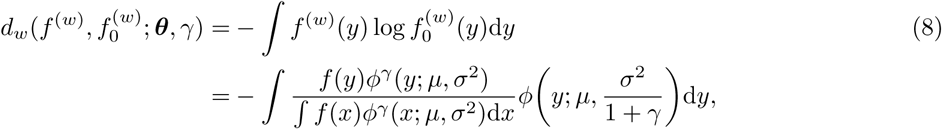

where *f* ^(*w*)^ and 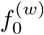 are defined in (5) (replacing ***θ***_0_ by ***θ***). The empirical cross entropy on the samples is

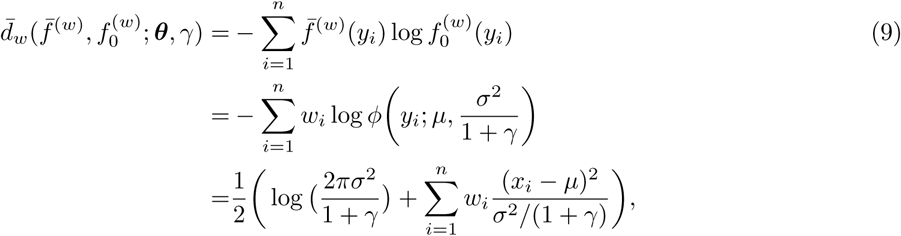

where *w*_*i*_ is defined in (7). Given a fixed *γ*, define

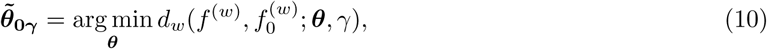

and its M-estimate

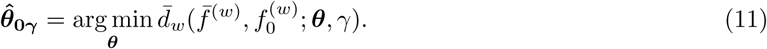

Since the weights reply on the parameter we would like to estimate, we can iteratively update the weights and ***θ***. The minimizer 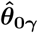 is the fixed point. From our practical experience and previous works, the convergence of the algorithm is not a problem. In the step of given the weights *w*_*i*_’s to update ***θ***, setting the derivative of 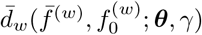 with respect to ***θ*** to be zero gives

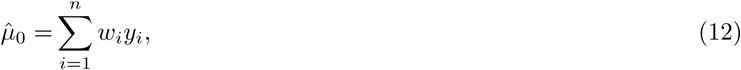

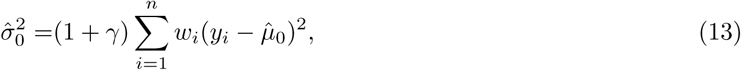

which is the exact estimation equation in (4). Rearranging the terms in (4) gives

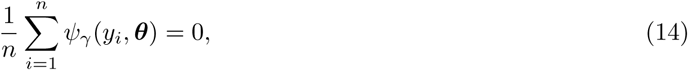

where 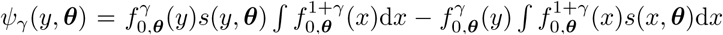 and 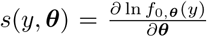, which is the estimating equation for *γ*-cross entropy (Fujisawa and Eguchi, 2008). Our M-estimate from the cross entropy criterion (9) agrees with the estimator from Windham’s procedure and *γ*-cross entropy criterion, and thus they share the same consistency property and asymptotic normality property stated in Proposition 1. In the case that underlying density is purely Gaussian without outliers (*π*_1_ = 0), the asymptotic variances for 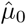 and 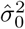 are

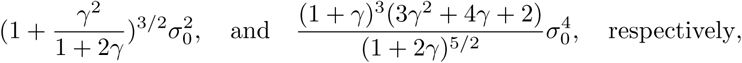

also shown in (Basu et al., 1998; Jones et al., 2001). From Proposition 1, setting *γ* = 0 gives that the asymptotic variance for 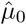 is 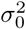 and for 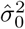 is 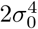, which are the most efficient asymptotic variances for MLEs. If the predefined *γ >* 0, their asymptotic variances increase with *γ*.

**Proposition 1**. *If there is no outlier that* 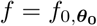, then 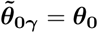 *and thus the estimate is Fisher consistent. Under the mild conditions, applying the theorems for M-estimate in Van der Vaart (2000), shown in Fujisawa (2013); Fujisawa and Eguchi (2008)*,

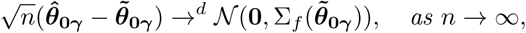

*where* 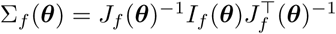, 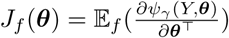 *and* 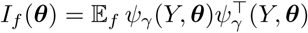.

The population proportion can be simply estimated from the view of weighting the mixture model. Consider taking the integral of unstandardized density-power-weighted mixture model,

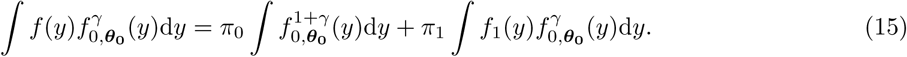

Rearranging (15) gives

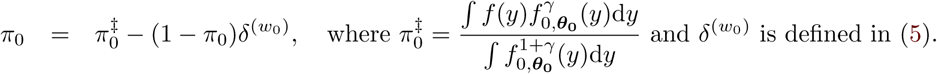

Hence, an upward bias estimator for *π*_0_ is

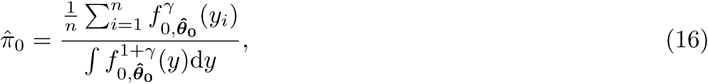

which agrees with the result in Kanamori and Fujisawa (2015) by minimizing their unnormalized density power score.

### 2.4 Bias and variance trade-off in *γ*

So far we have seen that the M-estimate 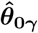 and its asymptotic limit 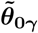 depend on a predefined *γ*. Proposition 1 shows that 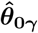 is *√n*-consistent to 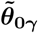. However, there is still a latent bias from 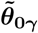 to the target parameter ***θ***_0_, also pointed out in Fujisawa (2013). The weighted mixture model (5) is the ideal model where the ***θ*** in the weighting density 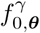 is the underlying ***θ***_0_. However, actually the weighting density we fit is under 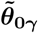 (supposed we have an infinite amount of samples), and the fitted weighted mixture density is

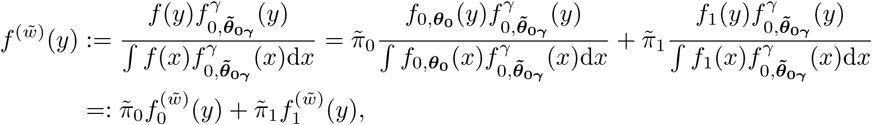

where 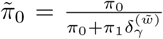, 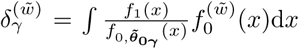 and 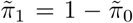, which are in the similar notations as in (5) except the superscript is changed from (*w*_0_) to (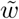). Consider the Gaussian mixture model, 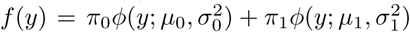. The asymptotic estimates for (*µ*_0_, 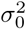) from minimizing (8) are

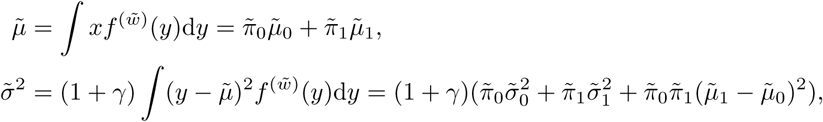

where 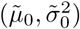 are the parameters in 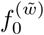 (density for 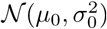) and 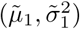 in 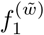 (density for 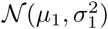). Their M-estimates are in (12) and (13). Based on some basic calculations,

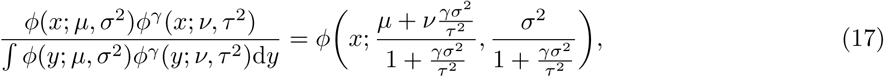

and

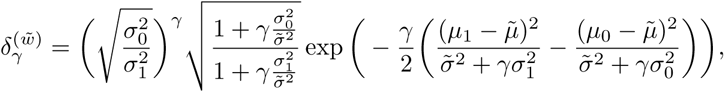

we get 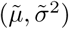 satisfying

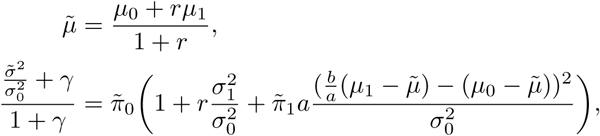

where 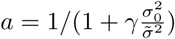, 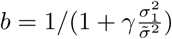, and

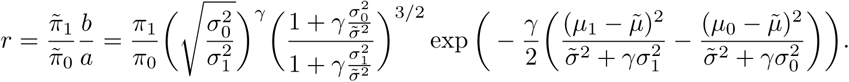

Suppose 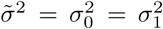 known. Increasing *γ* accelerates 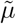 approaching to *µ*_0_. The form of 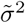 is much complicated, which depends not only on *γ* but also how close of 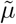 to *µ*_0_ and the underlying parameters (*µ*_0_, *µ*_1_, 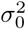, 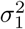, *π*_0_). Hence, the latent bias is model dependent, relying on the choice of *γ* and usually hard to quantify.

From a classical point of view in robust estimation, we investigate the Influence Function (IF) to see how it varies under different *γ*’s. Previous works also studied the IF of density weight divergence in (Basu et al., 1998; Jones et al., 2001), but here we more focus on how it is affected by *γ*. The IF for normal mean estimate 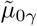 and the IF for normal variance estimate 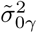 from *γ*-robustifying procedure at normal distribution 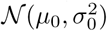 are

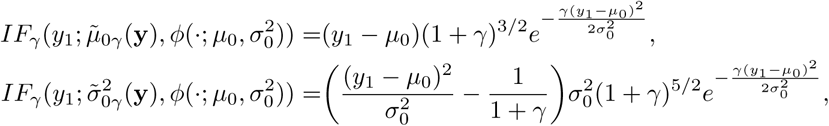

where 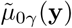 and 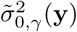 are the minimizers defined in (10). The IF provides us a heuristic tool considering if there are an infinitesimal amount of outliers lying at point *y*_1_, how the outliers affect the asymptotic bias of the estimate (Hampel et al., 2011). We demonstrate the IF curves at (0, 1) under various *γ*’s in Figure 2. When *γ* = 0, both IFs of the mean estimate and the variance estimate are unbounded, while when *γ >* 0, the IFs are redescending and thus the estimates are robust. In the case that the outlier *y*_1_ is more than 2 standard deviation away from the underlying mean zero, a larger *γ* has a better performance to down weight the outliers, whereas in the case that the outlier is near zero, a large *γ* can cause even bigger bias than that of MLE (corresponding to *γ* = 0). When *γ* ≥ 0.5, the maximums of both the gross error sensitivities of the IF of mean estimate and the IF of variance estimate under various *γ*’s in the comparison are at *γ* = 3. The IF approach only considers an infinitesimal amount of outliers, while in reality, there could be a heavy proportion of outliers in various magnitudes. Besides, there needs a balance to choose a *γ* to minimize the estimation error for the mean and also the error for the variance, which makes the problem of *γ* selection more difficult.

**Figure 2:**
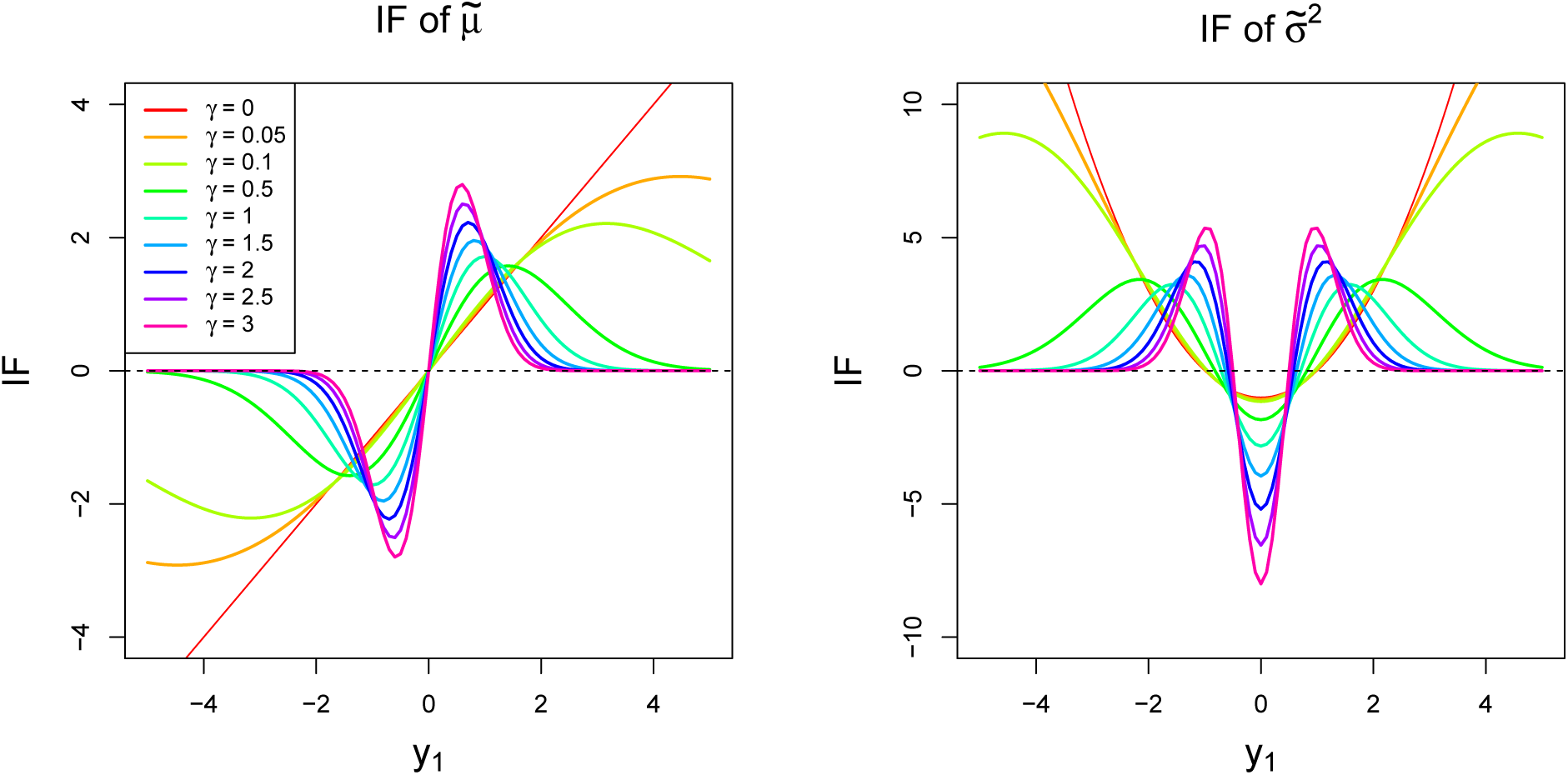
Influence functions under various *γ*’s of 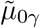 (in the left panel) and 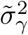 (in the right panel) at standard normal distribution 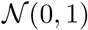.

We demonstrate how *γ* balances bias and variance of the estimation in simulation studies. Consider 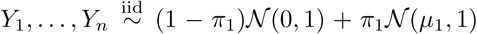, where *n* = 200, 2000 and under each *n*, consider (*µ, π*) = (0, 0), (1, 0.1), (3, 0.1), (5, 0.1), (1, 0.3), (3, 0.3), (5, 0.3). The population parameters are set as *µ*_0_ = 0, 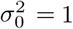. We evaluate the MSE for estimating *µ*_0_ and *σ*_0_ under *γ* = 0, 0.5, 1, 2, 3 from *B* = 50 realizations, i.e., 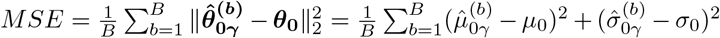, where the superscript (*b*) represents the *b^th^* realization. From Figure 3 — Figure 6, the MSE trends in the top row in each figure show that a too small *γ* leads a large bias and a too big *γ* leads large variance. Such trade-off phenomenon is very clear under *µ*_1_ = 3, 5 and *π*_1_ = 0.1, 0.3 especially under large sample size, while under cases such as *µ* = 1 and *π*_1_ = 0.1, 0.3, the MSE trends are almost flat. In the simulations, each realization can vary a lot and also is affected by sample size. The estimation trends of 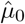, 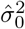 and 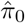 varying with *γ* are summarized in Figure S1 — Figure S4. From these simulation studies, we can see the performance of the estimation depends on *γ*, and the optimal *γ*, the turning point of the MSE curve, varies from case to case relying on the proportion and the magnitude of outliers and very data-dependent.

**Figure 3:**
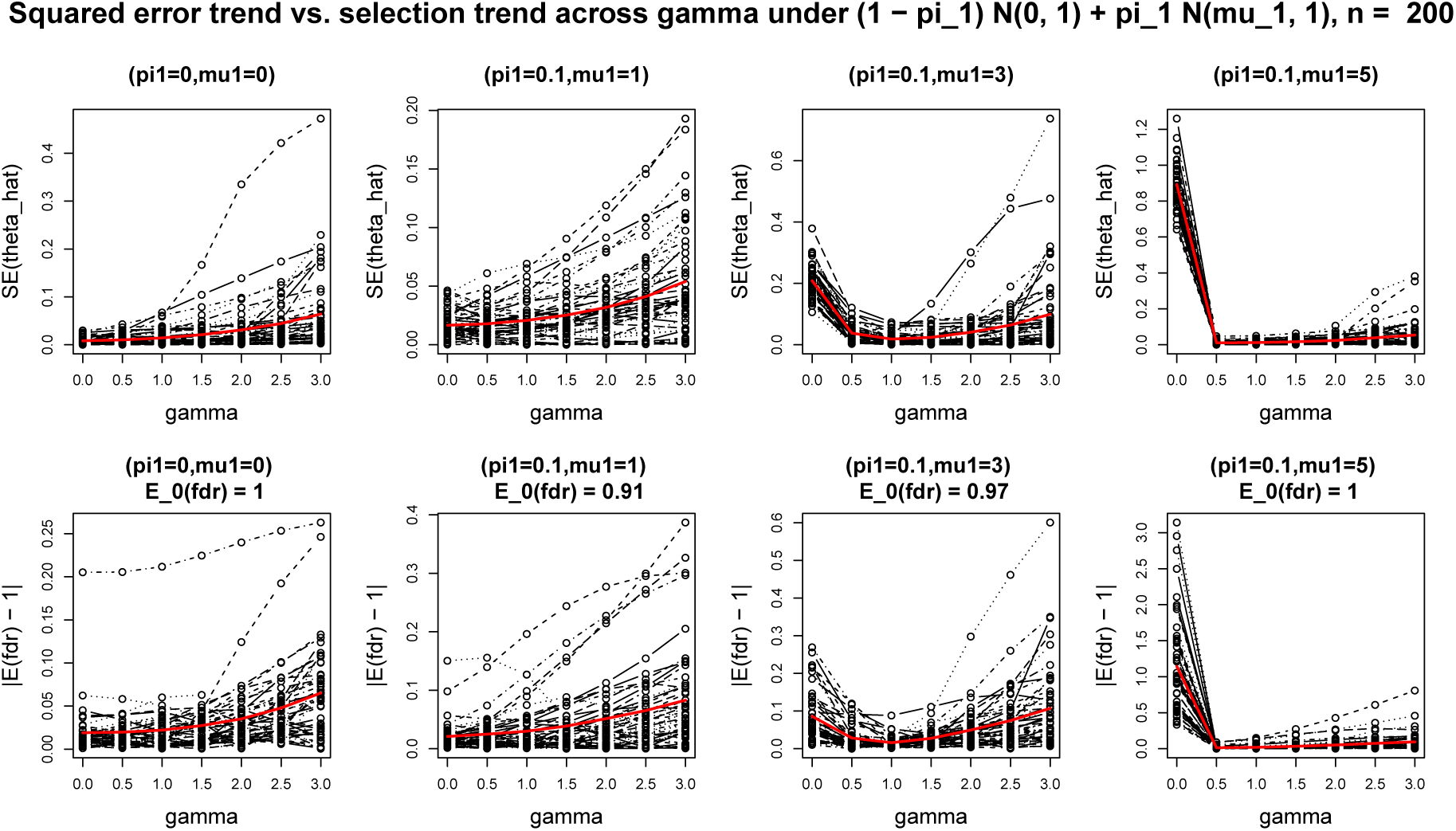
Comparison of squared error trend 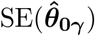 (in the top row) and selection trend 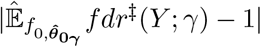 (in the bottom row) across *γ* = 0, 0.5; 1, 2, 3 under Gaussian mixture model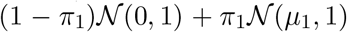 where *n* = 200 and (*μ*_1_, *π*_1_) = (0, 0), (1, 0,1); (3, 0.1), (5, 0.1). Each black curve is from one sample realization. The red curves in the top row are the MSE trends and in the bottom row are the average of the selection trends from 50 black curves at each *γ*. The oracle 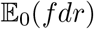 is indicated in the subtitle for each case.

### 2.5 Selection criterion on *γ*

Windham (1995) provided a selection method for *γ* from a view of asymptotic efficiency, relating the convergence rate of the iterative algorithm to the asymptotic variance. However, it did not consider the latent bias from the true population parameter. The analysis in Fujisawa and Eguchi (2008) is based on the assumption that under some *γ*, the outliers lie in the tails of the population density raised to power *γ*. Under this assumption, they claimed the latent bias is small. However, this assumption is not guaranteed for each *γ*. As far as we know, how to select a “proper” *γ* is still unknown for the mixture model.

Recall that the *γ*-robustifying procedure actually approximates *f* ^(*w*)^ to be 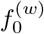. What we expect is to weight more on the population points while less on the outliers. However, only letting *f* ^(*w*)^ close to 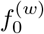 does not guarantee that this approximation is overall good for the population points. As we have seen in Subsection 2.4, there is a bias-variance trade-off controlled by *γ*. Since we are mainly interested in estimating the distribution of the population points, we consider a goodness-of-fit (GOF) measurement on how close of 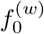 to *f* ^(*w*)^ in the probability measure of the population distribution and thus give an overall GOF on the population points,

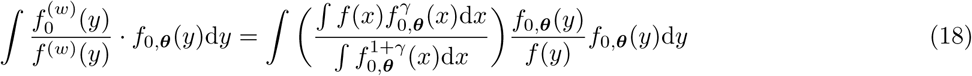

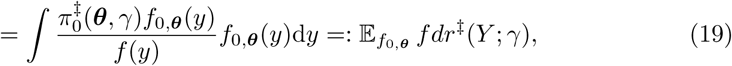

where 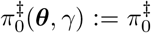 defined in (16) and 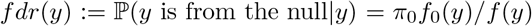 is the local false discovery rate introduced in Efron (2005). Here we replace the true *π*_0_ by 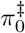 and add the same superscript ‡ on *fdr*. The oracle value for 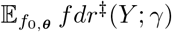 is the ordinary expected fdr under the null, i.e., 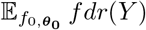 when 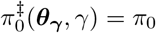 and ***θ***_***γ***_ = ***θ***_0_.

Consider under some *γ*, 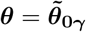 which is the asymptotic limit of the M-estimate. We have 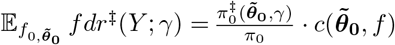, where

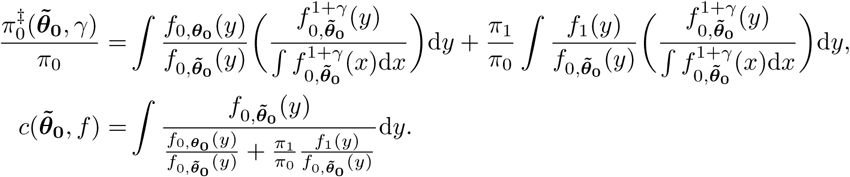

If the *γ* is well selected such that 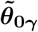 close to ***θ***_0_ and 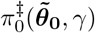 close to *π*_0_, then 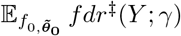 will be close to the ordinary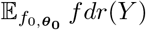. For example, under the Gaussian mixture model 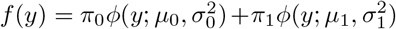, we have

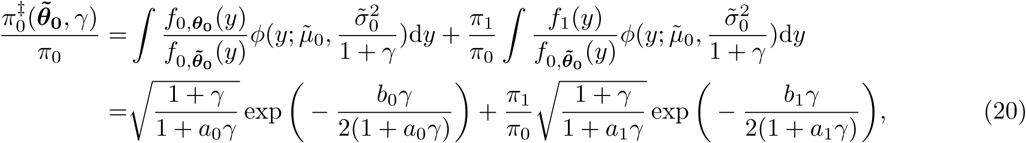

where 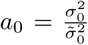, 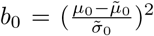, 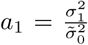, and 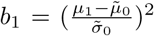. Under a proper *γ*, 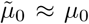 and 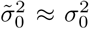 so that *a*_0_ *≈* 1 and *b*_0_ *≈* 0, thus the first term in the Right Hand Side (RHS) of (20) is close to 1. Further if *π*_1_ is much smaller than *π*_0_ or 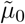 is far away from *µ*_1_ relative to the scale 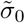, i.e., *b*_1_ is large, then the second term of the RHS of (20) is close to 0. Hence, under such *γ*, 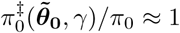 and 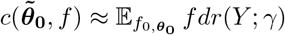. In a special case that *π*_1_ = 0, by Fisher consistency of 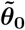 from Proposition 1, for all *γ ≥* 0, 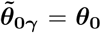 and thus 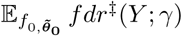 reaches to its oracle 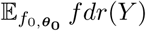, which is one in this case. In reality, since we do not know the underlying mixture model, we would like to approximate the oracle 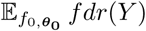 to be one, although this approximation may not be accurate in some cases where *f*_0_ and *f*_1_ are close to each other and the outlier proportion is not small. By comparing 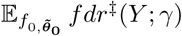 to one tries to capture the latent bias hidden in 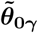.

To estimate 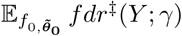 in practice, since the expectation of *fdr* is with respect to the population, here we do not care too much of the tail effect when estimating the density functions. We discretize the integral into bins and estimate the density by its histogram in each bin. In details, we first partition the range of the observed *y*’s to *S* non-overlap intervals, i.e., 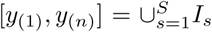 where *y*_(*i*)_ is the *i*^th^ smallest value, and *I*_*s*_ ∩ *I*_*t*_ = ∅ for *s* ≠ *t*. Define |*I*_*s*_| ≠ #{*y*: *y* ∈ *I*_*s*_}. We then merge the adjacent bins such that for all |*I*_*k*_| strictly positive. In one bin *I*_*k*_, we approximate the integral of mixture density *f* over *I*_*s*_ by sample proportion 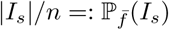 and the integral of *f*_0_ over *I*_*s*_ is 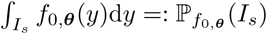. For the most left and right intervals, we extend them to −∞ and +∞, respectively, when calculating 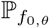. Finally, we get an empirical estimate for 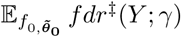 from

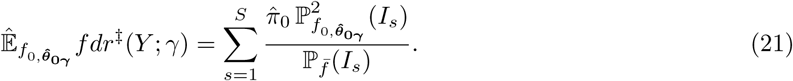

Since this empirical estimate (21) depends on 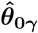, this term tries to capture the variation of 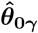 under different *γ*’s.

Our selection criterion is to select a *γ* minimizing the distance between the empirical estimate for 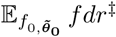 and the approximation one for the oracle expectation, i.e.,

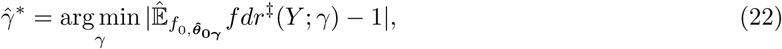

where 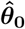 is our robust estimate from (11). In the ideal case, the optimal *γ*^∗^ is the minimizer of 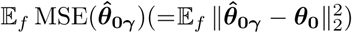. But it is impractical since MSE depends on unknown ***θ***_0_. Our contribution is to transfer the inestimable criterion to an estimable criterion in order to select a proper *γ* in practice. Observe that

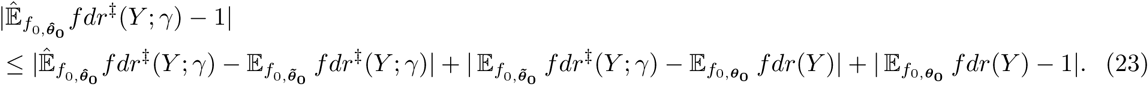

We expect around the optimal *γ*^∗^, the first term in the RHS of (23), which controls the variance, is increasing with *γ*, and the second term in the RHS, which controls the bias, is decreasing with *γ*. Hence, there would be a turning point, which indicates the occurring of a proper *γ*. The third term in the RHS is the unavoidable bias from the selection procedure. Since the estimate in (21) not only depends on the estimate for ***θ***_0_ and *π*_0_, but also on the estimate for the density ratio, due to its complicated form, we study the behavior of our selection trend via simulations. In the same setting as in the simulations in Subsection 2.4, we compare the inestimable Squared Error (SE) trend 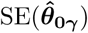 versus our practical selection trend 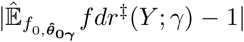 across various *γ*’s in Figure 3 – Figure 6. Overall, comparing the average performances, our selection trends have similar capability in average performance as the MSE trends to capture the minimizer *γ* under various settings and under both small and large sample sizes. In the case that there is no outlier (*µ*_1_ = 0, *π*_1_ = 0), both MSE trend and the average selection trend increase in *γ*. In the case of well-separated mixture densities under (*µ*_1_ = 5, *π*_1_ = 0.1), there is a clear turning point at *γ* = 1. In the case of hardly separated mixture densities under (*µ*_1_ = 1, *π*_1_ = 0.3), where the oracle 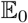 *fdr* is around 0.76, although approximating 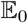 *fdr* to be 1 is not quite accurate, the SE trends and our selection trends are almost flat in average. In the cases of moderately hard separated mixture densities under (*µ*_1_ = 3, 5, *π*_1_ = 0.3), our average selection trend can have two local minimums, one in a large *γ* and another one near *γ* = 0. When we look at the 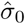 under different *γ*’s in Figure S2 and Figure S4, the trends first increase from *γ* = 0 then decrease, indicating our selection trends are more likely affected by 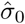 in these cases. And under (*µ*_1_ = 3, *π*_1_ = 0.3) especially in small sample size, the realization curves have large variations.

**Figure 4:**
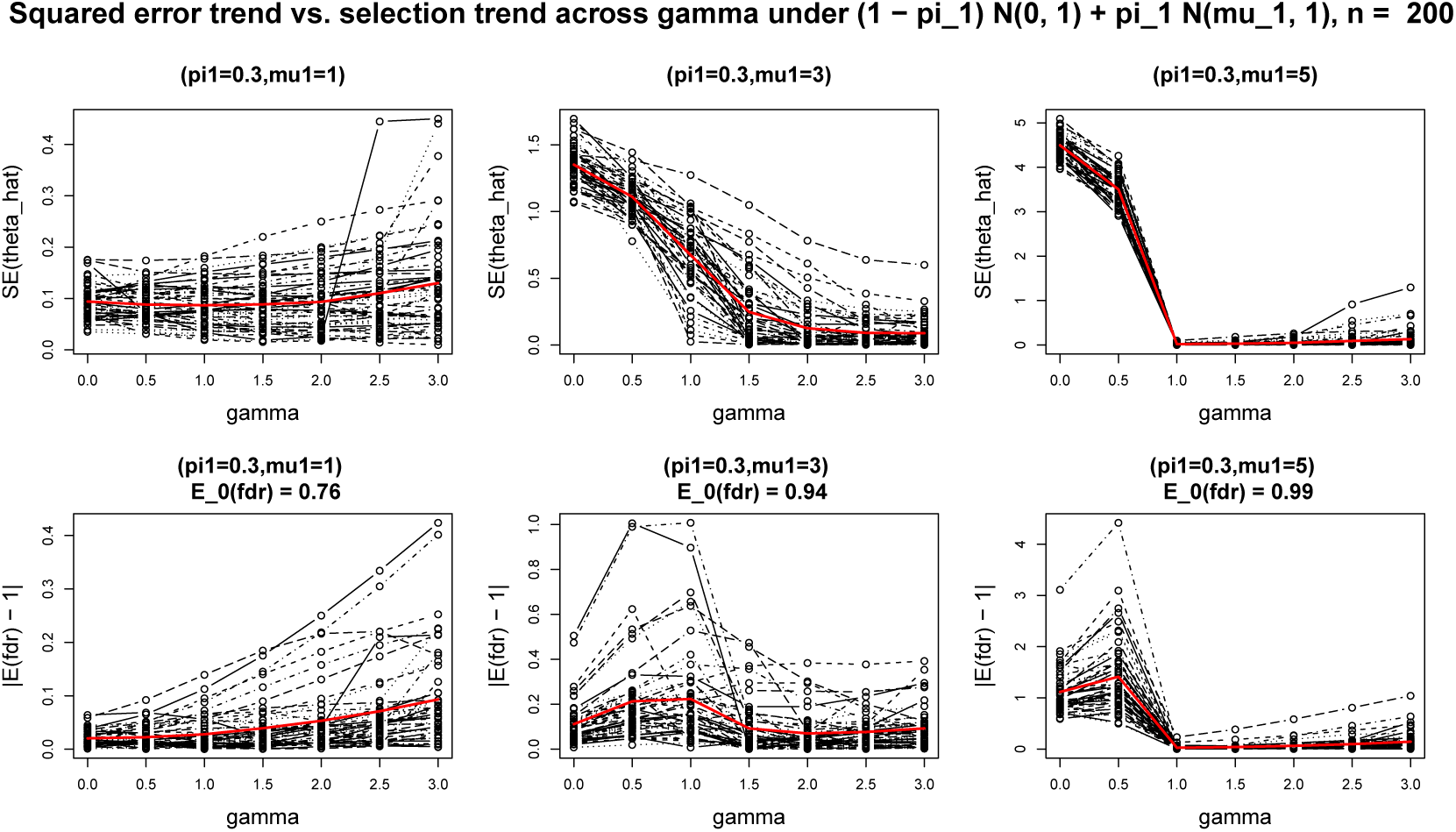
The same caption as Figure 3 under *n* = 200, (*μ*_1_, *π*_1_) = (1, 0.3), (3, 0.3); (5, 0.3).

**Figure 5:**
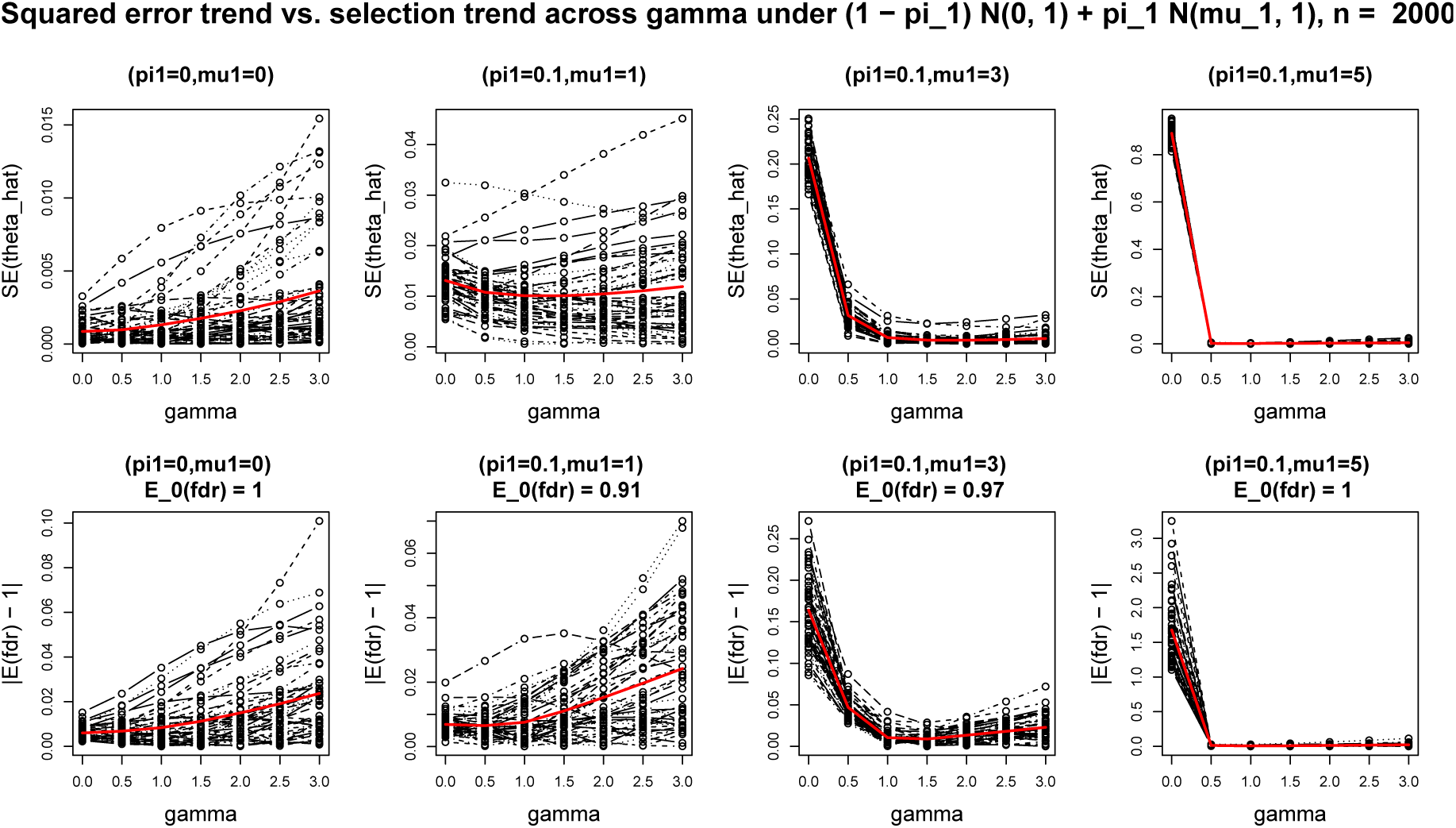
The same caption as Figure 3 under *n* = 2000, (*μ*_1_, *π*_1_) = (0, 0), (1, 0.1); (3, 0.1); (5, 0.1).

**Figure 6:**
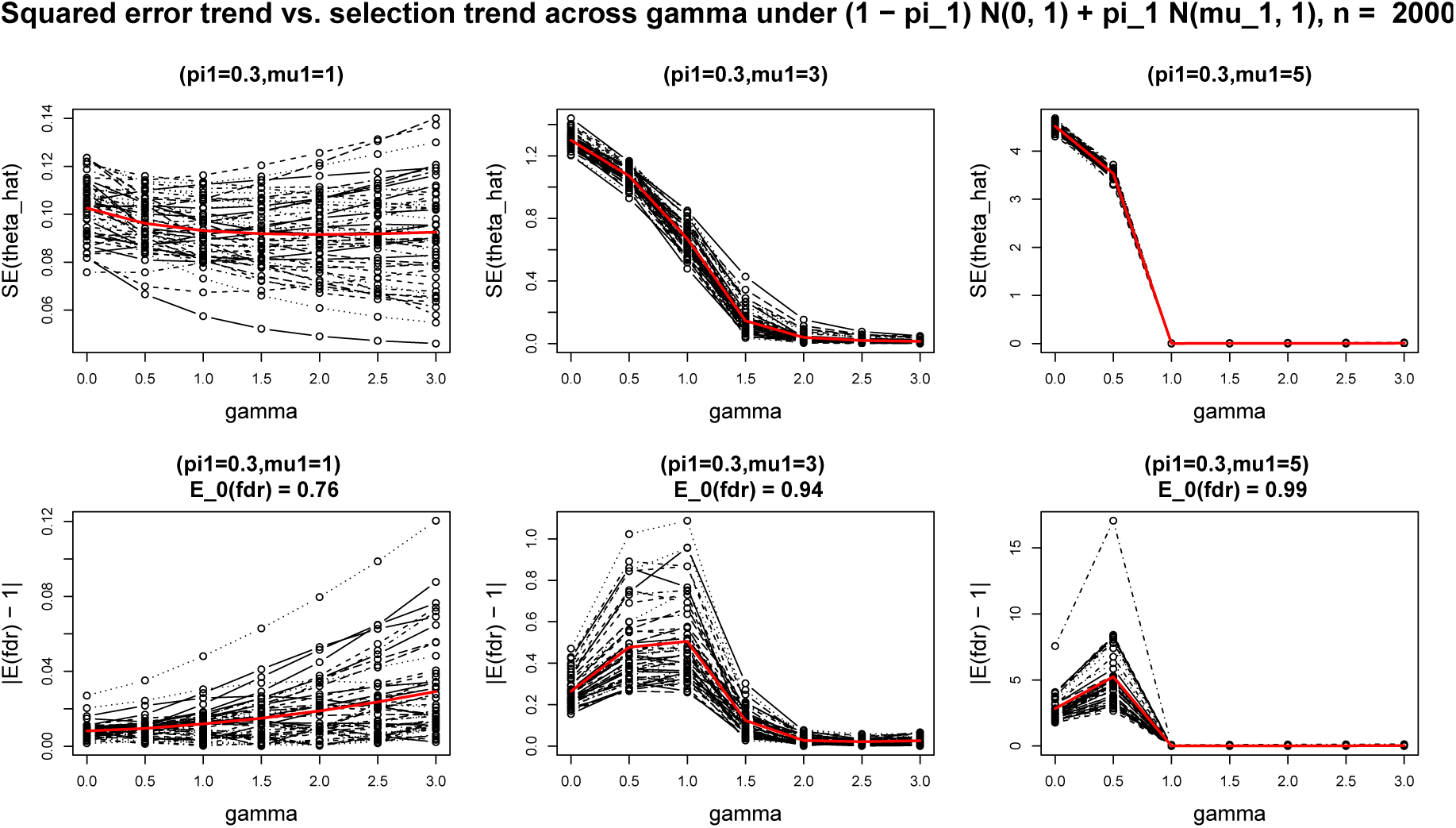
The same caption as Figure 3 under *n* = 2000, (*μ*_1_, *π*_1_) = (1, 0.3), (3, 0.3), (5, 0.3).

(Huber, 2011) pointed that the redescending M-estimate could have multiple minima and can be locally trapped. When there are multiple comparable and well-separated density bumps, our selection may pick up the wrong bump as the population especially in the small sample size, but such case is easy to detect. In the real application, we add one diagnosis step on the fitted residuals to flag the possible problematic fitting. We detail it in Subsection 2.6.

### 2.6 Data-adaptive algorithm in linear regression

So far, we consider the response variable **y** only from one group, mainly affected by one covariate variable as in (3), i.e.,

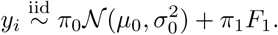

Now we go back to the general setting in linear regression (1)–(2), i.e.,

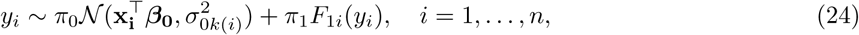

We still assume independence among *y*_*i*_’s but now *y*_*i*_ comes from different populations since 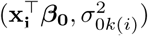 varies with *i*. Hence, we need first to construct the null samples from the same distribution by transforming *y*_*i*_ to a standardized expression, Consider the standardized residual for the *i*^*th*^ sample,

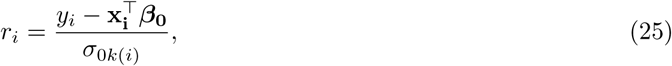

where 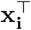 is the *i*^*th*^ row of matrix **X** and the index *k*(*i*) indicates the group of the *i*^*th*^ sample. Under the mixture model of **y**, the standardized residuals are i.i.d. from the mixture model

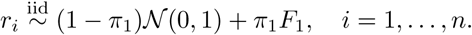

To get empirical cross entropy between 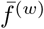 and 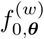 defined in (9) on sample *y*_*i*_’s, by changing variables from the residuals to **y**, we have

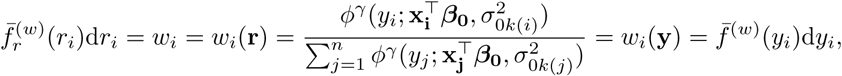

and

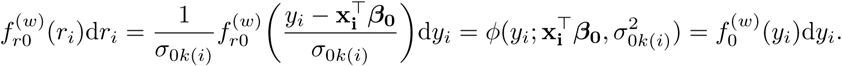

Hence, in linear regression problem, our robust likelihood criterion in terms of cross entropy on the weighted samples is

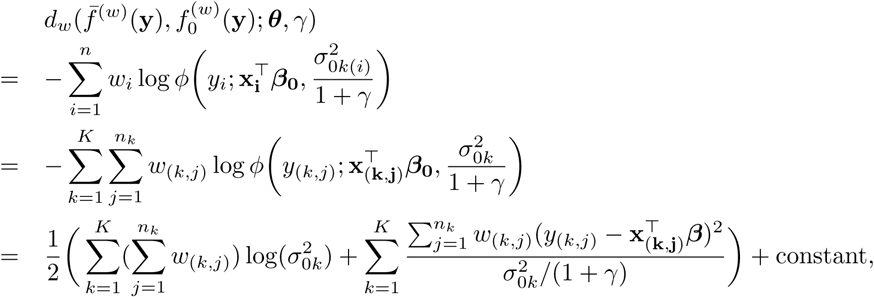

where the pair index (*k, j*) indicates the *j*^*th*^ sample in group *k*, *k* = 1, …, *K* and *j* = 1, …, *n*_*K*_. Therefore, given *γ*, the M-estimate for *θ*_**0**_ is

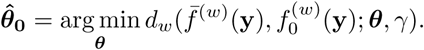

The minimizer can be found by iterating the weights and the estimate for ***θ***_0_. After updating the standardized residuals based on the estimate for ***θ***_0_ in each iteration, we apply our data-adaptive *γ* selection procedure of **Algorithm 1** to select a proper *γ* to further optimize the weights. The whole procedure is summarized in **Algorithm 2**.

## 3 Simulation Studies and real data application

In this section, we apply our algorithm AdaReg in the simulation studies to compare with other robust procedures then apply it in the real dataset of heart samples from RNA-seq in GTEx project.

### 3.1 Comparisons of robust procedures in simulation studies

In the simulation studies, we compare the performances of several robust estimation methods in a series of settings to evaluate their estimation accuracy. The methods under comparisons include the OLS (*γ* = 0), *γ*robustifying methods under *γ* = 0.5, 1, 2, 3 and our data-adaptive robustifying procedure (AdaReg), other Mestimation methods based on Huber’s weight (Huber, 2011), Hampel’s weight (Hampel et al., 2011), Tukey’s bisquare weight, and S-estimation (Rousseeuw and Yohai, 1984), and resistant robust methods including the least median squares (lms) (Rousseeuw, 1984), and the least trimmed squares (lts) (Rousseeuw, 1984, 1985). The OLS is implemented by function *lm*(*⋅*) in R (Team et al., 2013). The fixed *γ* and data-adaptive robustifying procedures are from **Algorithm 1-2**. The other M-estimations are from function *rlm*(*⋅*) in package “MASS” (Venables and Ripley, 2013). The lms, lts and S estimation are from function *lqs*(⋅) in package “WRS2” (Mair and Wilcox, 2016).

#### Algorithm 1: Data-adaptive *γ* selection procedure

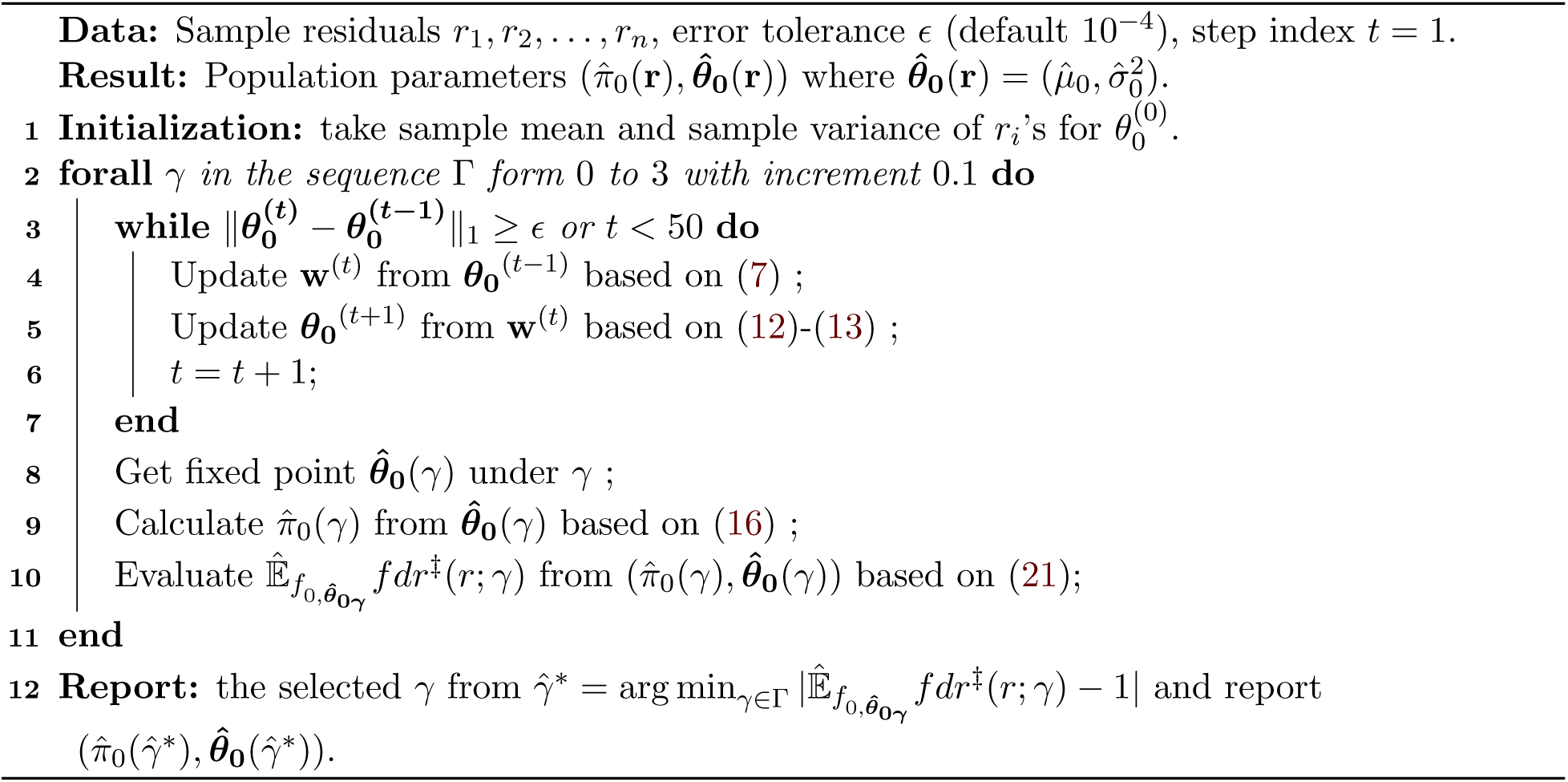

#### Algorithm 2: Data-adaptive robust estimation in linear regression

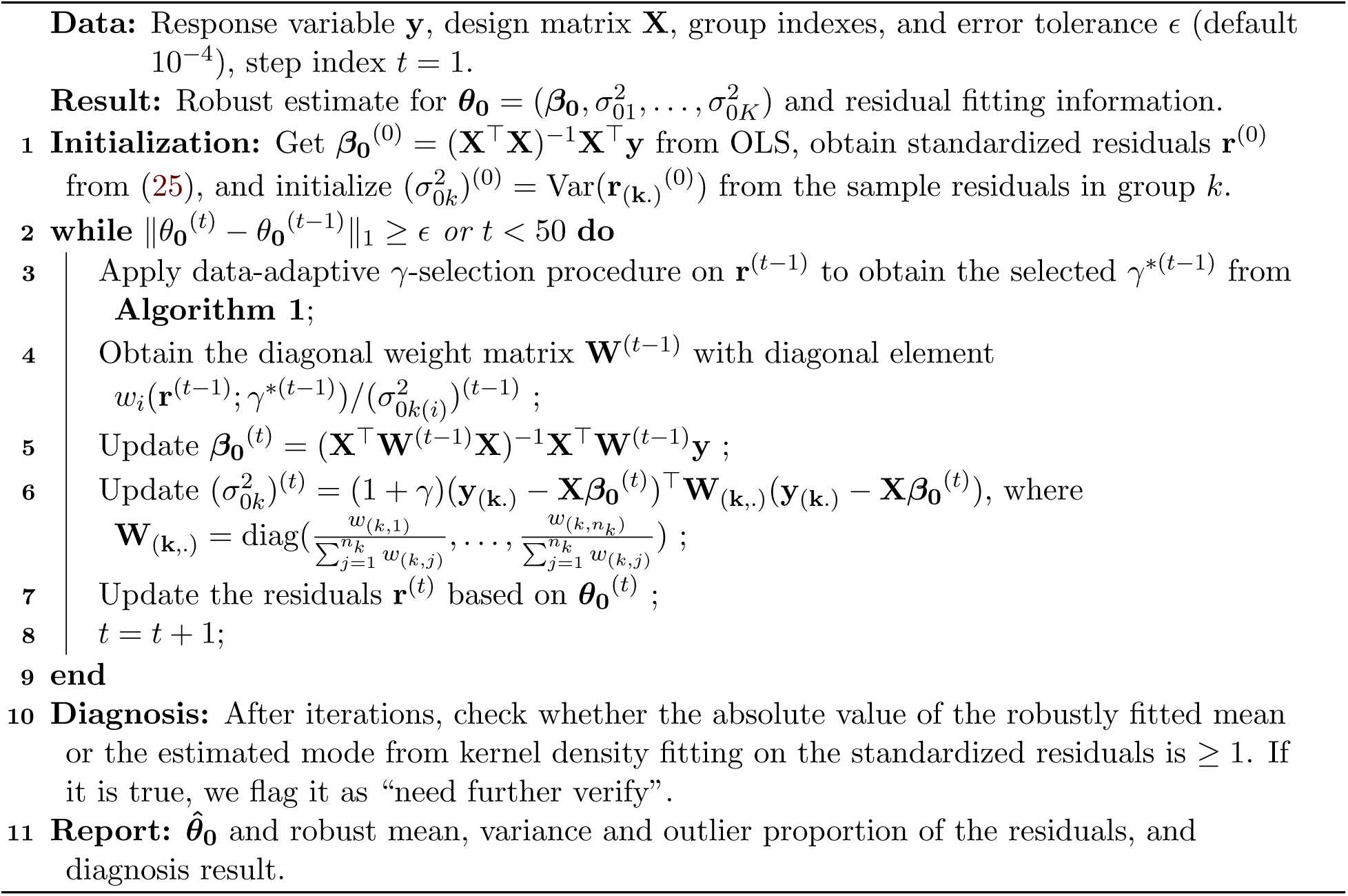

We consider three cases: (i) a small sample size and small dimension case of *n* = 200 and *p* = 2 (including the intercept), (ii) a moderately large sample size and small dimension case of *n* = 2000 and *p* = 2, and (iii) a moderately large sample size and moderately large dimension case of *n* = 2000 and *p* = 20. In each case, we set ***β***_0_ = **1**_**p×1**_. For the design matrix **X**, its first column is 1’s and we generate its other elements i.i.d. from 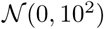. We consider the elements of the mixture noise ***ϵ*** i.i.d. from 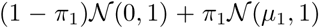 and set (*π*_1_, *µ*_1_) from the grid (0.1, 0.2, …, 0.4) (1, 2, …, 10) and (0, 0). In each case, we evaluate the square root of the mean square error (RMSE) for ***β***_0_ and 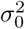 for each method from *B* = 100 independent repeated procedures, where 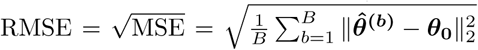. The results are summarized in Figure S5–Figure S6 for case (i) (*n* = 200, *p* = 2), Figure S8–Figure S9 for case (ii) (*n* = 2000, *p* = 2), Figure S11–Figure S12 for case (iii) (*n* = 2000, *p* = 20). We further investigate method detection ability when *π*_1_ > 0 by evaluating the True Positive Rate (TPR, the proportion of the rejections over the true positives) and the False Positive Rate (FPR, the proportion of the non-rejections over the true negatives). We reject a sample if its standardized residual score 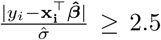. We report the average TPR and average FPR for each method under each case from 100 independent repeated procedures summarized in Figure S7 for case (i) (*n* = 200, *p* = 2), Figure S10 for case (ii) (*n* = 2000, *p* = 2), and Figure S13 for case (iii) (*n* = 2000, *p* = 20). For the cases of large proportion of outliers under *π*_1_ = 0.4 and large *µ*_1_ ≥ 3, it could happen that our algorithm picks the wrong bump as the population distribution, especially in small sample size as mentioned in Subsection 2.5. Hence, we filter the cases that the absolute value of the fitted mean for the standardized residuals ≥ 1 for AdaReg in the comparisons under the setting of *n* = 200, in total less than 20 realizations in the simulations.

To compare the estimations for ***β***_0_ from Figure S5 (*n* = 200, *p* = 2), Figure S8 (*n* = 2000, *p* = 2) and Figure S11 (*n* = 2000, *p* = 20), in each case we can see the fixed *γ*-robustifying procedure from *γ* = 0 (OLS) to *γ* = 3 gradually perform better and better, that is, the high RMSE region is smaller and smaller. The Huber and Hampel weighted robust methods perform worse than the *γ*-robustifying procedures. The Tukey’s bisquare estimation has similar performance as the *γ* = 0.5 robustifying procedure. The Rousseeuw’s methods (lms, lts, S estimation) perform similar to the large *γ* ≥ 2 robustifying procedure in the low dimension case (*p* = 2) but become worse in the case of moderately high dimension case (*p* = 20). Our AdaReg performs better than the fixed *γ*-procedure overall, slightly worse in the situations of the null and the alternative hardly separable such as in (*π*_1_, *µ*_1_) = (0.3, 2), (0.4, 3). To compare the convergence rate, the small *γ*(≤1) procedures and Huber, Tukey and Hampel’s estimations have similar performances from case (*n* = 200, *p* = 2) to (*n* = 2000, *p* = 2), respectively, while for the large *γ*(*>* 1) and data-adaptive robustifying procedures have better performances when sample size increases, which indicates that their estimation convergence rates are relatively slow. To compare the dimension effect of the explanatory variables from case (*n* = 2000, *p* = 2) to (*n* = 2000, *p* = 20), the number of explanatory variables does not affect much for *γ*-robustifying procedures, Huber, Tukey and Hampel methods, since they down weight the outliers in the residuals which is essentially in one dimension. Although the S-estimator performs well in low dimension, it loses its advantage in high dimension. However, for Rousseeuw’s methods (lms, lts), they search the population points in the (1 + *p*)dimensional space, and thus high dimension dramatically diminishes their performances.

We also compare the estimations for population variance 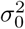 of the noise and outlier detection abilities, summarized in Figure S6–Figure S7 (for *n* = 200, *p* = 2), Figure S9–Figure S10 (for *n* = 2000, *p* = 2) and Figure S12–Figure S13 (for *n* = 2000, *p* = 20). To robustly estimate the population variance fo the noise is important for the outlier detection. An upwards biased estimate can lead high false negative rate and a downwards biased estimate can have high false positive rate. From the simulations, the *γ*(*>* 0) robustitfying procedures perform better than the traditional robust estimation methods to estimate *σ*_0_. In the case of (*n* = 2000, *p* = 20), when *γ* = 3, the variance estimator is more likely to be affected by the robustness of 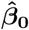 in high dimensions. For a fixed *γ*-robustifying procedure, it can be easily locally trapped. Our AdaReg shows its advantage in all cases to estimate the residual variance. From the plots of FPR versus TPR, our AdaReg preserves low FPR well in the hard detectable cases when *µ*_1_ *≤* 3 and achieves high TPR in the relatively easily detectable cases when *µ*_1_ *>* 3 over the other methods, no matter the sample size is large or small, the dimension is high or low, in our simulation studies.

### 3.2 Real Data Application in GTEx

We apply our AdaReg on the human gene expression data from RNA-seq in the GTEx project (Consortium et al., 2015) in version 7 (data downloaded from GTEx Portal). There are in total 600 heart samples from atrial appendage (297 samples) and left ventricle (303 samples). We consider the protein-coding genes whose either atrial sample median or left ventricle sample median is *>* 1 inTPM, in total 12, 422 genes. Here, we take logarithm transformation on (TPM + 1) making the data more symmetric.

We assume the expressions for each gene are independently from a mixture model 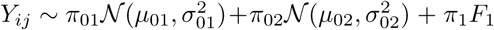, *j* = 1, …, 600, where *π*_1_ = 1 *− π*_01_ *− π*_02_. The population distribution for heart atrial appendage is 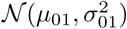 and for heart left ventricle is 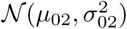. The outlier proportion is *π*_1_ and the outliers come from unknown distribution *F*_1_. We are interested in estimating (*µ*_01_, 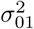, *µ*_02_, 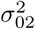, *π*_1_) under the presence of outliers in the gene expressions. We apply a series *γ*-robustifying procedures under *γ* = 0, 0.5, 1, 2, 3 and our data-adaptive procedure. Since from the simulation studies, the other methods cannot adapt to various settings, we do not compare them in this subsection.

Figure 7 shows the densities of 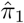 from the series *γ*-robustifying procedures under *γ* = 0.5, 1, 2, 3 and data-adaptively selected *γ*. Under *γ* = 0, which is OLS, since in this method 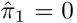, we do not show it in the density of plot of 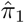. We can see under *γ* = 0.5, the outlier proportions are more concentrated near 0, while under *γ* = 2, 3, the procedures claim more outliers. The density of 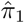 from AdaReg is comparable to the part of the density under *γ* = 1 in the estimated small outlier proportion region then approaches to the tail of the density under *γ* = 2 in the estimated high outlier proportion region, which indicates our data-adaptive procedure really tries to adapt to different outlier scenarios.

**Figure 7:**
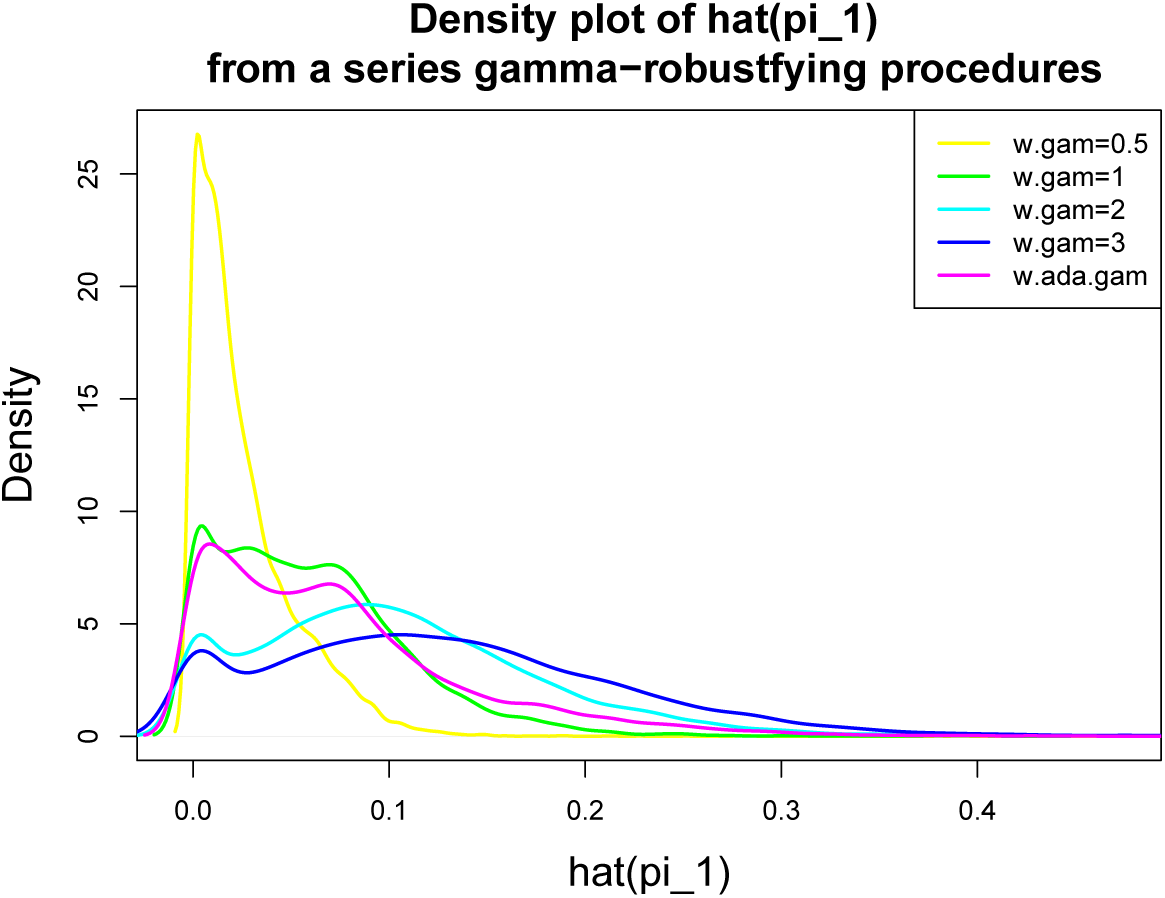
Density plot of 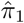 from a series *γ*-robustifying procedures under *γ* = 0.5, 1, 2, 3 and data-adaptive *γ* applied in heart expression data from GTEx.

In practice, it may be hard to check normality of the residuals without knowing which expression is an outlier. We here consider local property of the residuals. We estimate the density mode from kernel density fitting on the residuals for each estimation procedure. Suppose a procedure can estimate the population distribution well in each heart group then the standardized residuals should have a density peak around zero, as in the illustration example for gene MYH7 in Section 1. Figure 8 shows the scatter plot of 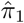 versus estimated standardized residual mode under each *γ*-robustifying procedure. Under *γ* = 0, 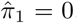, where we extend the values of the modes to a band in the figure. We can see under small *γ* = 0, 0.5, 1, there are a bunch of genes having positive residual modes away from zero, which indicates there is still some information left in the residuals, while under *γ* = 2, 3 and data-adaptively selected *γ*, the residual modes are more symmetric around zero. The residual modes from AdaReg are more concentrated at zero in the estimated small outlier proportion region (the red region), the same as under *γ* = 0.5, 1, and it still gathers majority of the residual modes being around zero in the estimated large outlier proportion region, the same as under *γ* = 2, 3. We pick the genes in the sparse point region (the blue region in Figure 8) as unfitted points. They may come from the result of being locally trapped. We put them aside for further analysis. In the following differential expression analysis, we do not include those unfitted genes. Note that checking from the residual modes does not give information on the normality region, which is harder to evaluate without prior knowledge. Here we only give some criteria for the not-well-fitted point.

**Figure 8:**
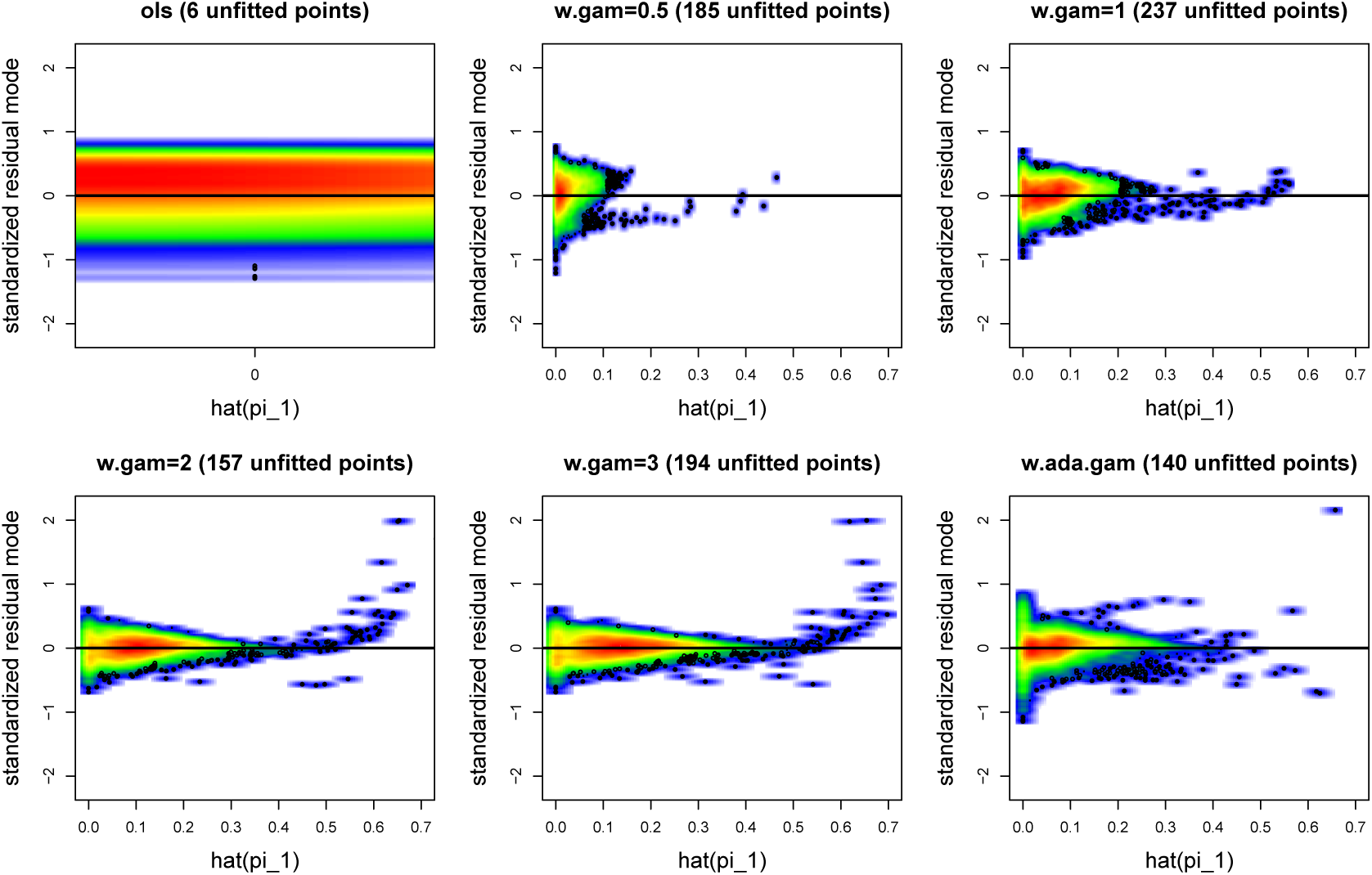
Scatter plot of 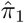 versus estimated standardized residual model from kernel density smoothing fitting under *γ*-robustifying procedures (*γ* = 0, 0.5, 1, 2, 3 and data-adaptive *γ*). The color from red to blue indicates the point density from dense to sparse. The black circled points in the blue region are defined as unfitted points. The horizontal black line is at zero.

We further investigate differentially expressed genes and enrichment analysis of GO terms (Consortium, 2014). From the *γ* robustifying procedures, we define the samples whose absolute values of the standardized residuals ≥ 2.5 as outliers then apply the two-sample *t*-test on the filtered samples after removing the outliers for each gene. Figure S14 shows the volcano plot of fold changes in log scale versus *p*-values from the *t*-test. We take the hyperbolic curve with curvature parameter 100 and minimum fold change parameter 1 as significance thresholds (Singh et al., 2016). The numbers of positive(negative) significant genes are comparable among all the procedures. In the GO term analysis from “clusterprofiler” R package (Yu et al., 2012), we combine the significant gene lists under *γ* = 0, 0.5, 1 as the results from small *γ* and combine the lists under *γ* = 2, 3 as the results from large *γ*. Figure 9 and Figure 10 show the significant enriched GO terms under threshold 10^*−*4^ on the adjusted *p*-values from the Benjamini-Hochberg procedure (Benjamini and Hochberg, 1995). For the genes highly expressed in left ventricle than in atrial appendage, they are highly enriched in ventricular cardiac muscle tissue functions, as expected. There the *p*-values from our AdaReg are comparable to the procedures under large *γ* and they are more significant than the results under small *γ* procedures. For the genes highly expressed in atrial appendage than in left ventricle, they are highly enriched in extracellular related functions. Again, the results from our AdaReg show the most significance.

**Figure 9:**
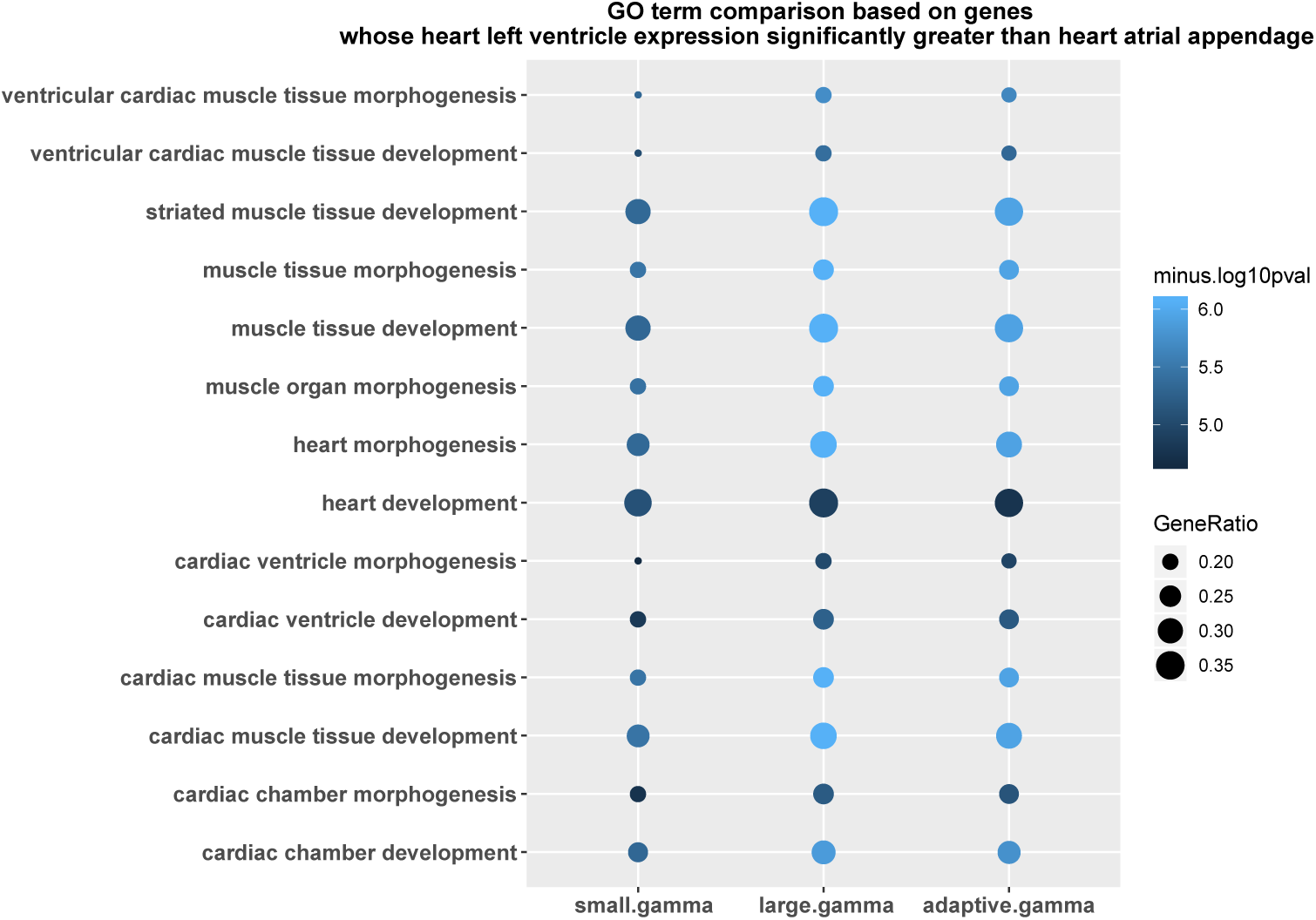
Bubble chart of minus of log_10_(*p*-value) for the significantly enriched GO terms (threshold: adjust *p*-values ≤ 10^−4^) under various *γ*-robustifying procedures including small *γ* (*γ* = 0, 0.5, 1), large *γ* = 2, 3 and our data-adaptively selected γ, based on the genes whose heart left ventricle expression significantly greater than heart atrial appendage expression.

**Figure 10:**
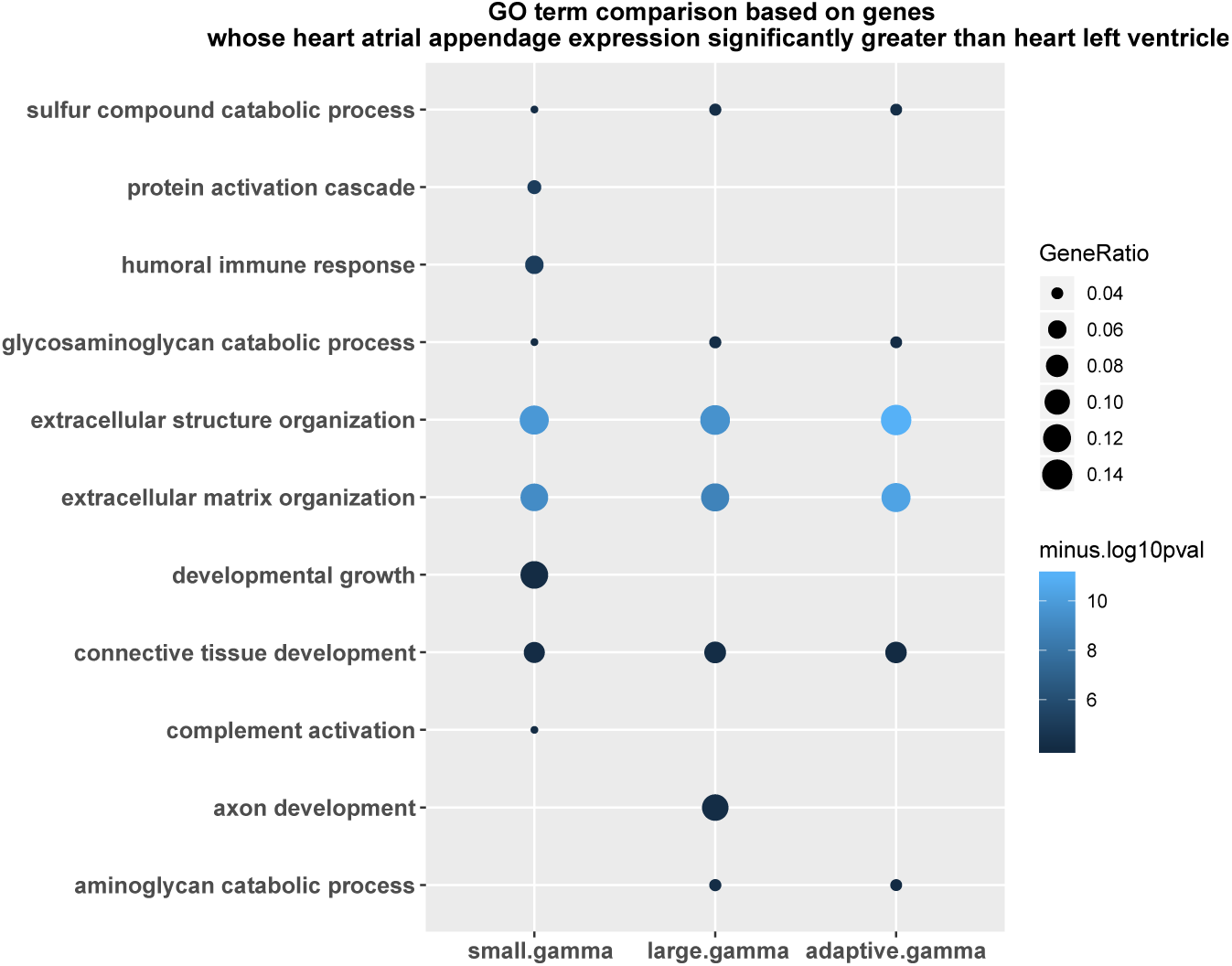
Bubble chart based on the genes whose heart atrial appendage expression significantly greater than heart left ventricle expression.

## 4 Discussion and conclusion

In the large-scale analysis of gene expressions, outliers vary from gene to gene. Given the presence of heterogenous outliers, it is important to have a robust estimation method to adapt to various genes. We followed the approach of the robust estimation based on *γ*-density-power-weight. Taking advantage that its estimation is only controlled by one tuning parameter *γ*, we developed a novel *γ* selection procedure to achieve the goal of data-adaptive estimation. We provided a heuristic analysis on the selection criterion, and found that our selection trends under various *γ*’s have similar capability to capture minimizer *γ* in average performance as the MSE trends from our simulations under a series of settings. Our data-adaptive robustifying procedure shows its advantage in both simulation studies and real data application compared to the fixed *γ* procedure and other robust methods in the setting of linear regression. However, there are still some limitations and further work to do.

### Assumption of the population distribution

In our setting, we mainly focused on the Gaussian population distribution. It is easy to generalize to other distributions like Poisson distribution, or Gamma distribution for different applications. Our robust criterion and selection procedure can be adapted easily to other settings. However, the approach of density-power-weight robust estimation requires the parametric assumption on the population.

### Outliers in the design matrix

In our setting, we only considered the outliers in the response variable. If there are outliers in the design matrix, the density-power-weight-based methods will lose their power. However, the resistant methods like the least median squares are still robust.

### Theoretical analysis of the selection criterion

To analyze the performance of our selection criterion, we provided a heuristic analysis of the selection criterion and compared our selection trend to the trend of MSE under various of *γ*’s in the simulation studies. We do not give a theoretical error bound for now because of the complicated form of the criterion, which depends not only on the parameter estimates but also the estimate for the density ratio. We think the error bound, either determinant or probabilistic, still depends on the underlying distribution and needs more theoretical work in the future.

### Other Selection criteria

The classical model selection criteria like Akaike’s Information Criterion (AIC) and the Bayesian Information Criterion (BIC) require that their sample distribution is known and thus do not work in our mixture setting with unknown outlier distribution. One may think about selecting *γ* from the diagnosis on the residuals. However, if the population is fitted wrong, the residuals could mask the true signals and thus could not give a fair selection on *γ*. This paper provides one way to select *γ*, but we look forward to other researchers developing other procedures or criteria to select *γ*.

In conclusion, robust estimation based on *γ*-density-power-weight is an interesting and important approach. We proposed one data-adaptive robustifying procedure: automatically selecting a proper *γ*. We believe that combining density-power-weight robust estimation with a data-adaptive *γ* selection procedure will be applicable to more situations involving the detection of differential gene expression, and the identification of tissue-specific genes.

## Acknowledgments

The Genotype-Tissue Expression (GTEx) Project was supported by the Common Fund of the Office of the Director of the National Institutes of Health, and by NCI, NHGRI, NHLBI, NIDA, NIMH, and NINDS. The gene expression TPM data used for the analyses described in this manuscript were obtained from GTEx Portal in version 7. We acknowledge the discussions with Dr. Hua Tang at Stanford. We would like to thank the funding supports by GTEx grant (5U01HL13104203) and CEGS grant (2RM1HG00773506).

## Supplementary Materials

**Figure S1:**
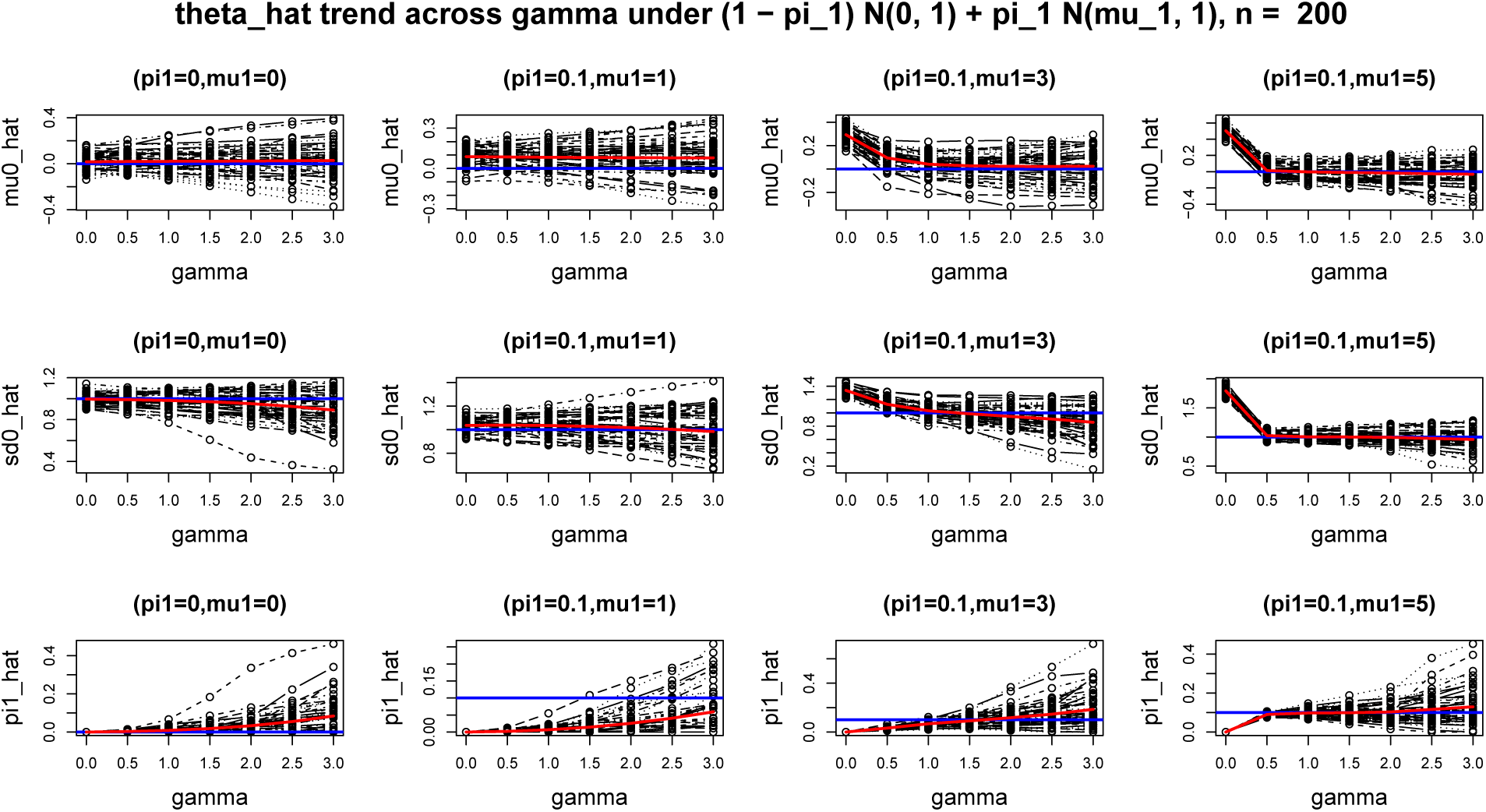
Trends of 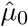, 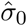 and 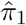 across *γ* = 0, 0.5, 1, 2, 3 under Gaussian mixture model 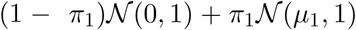 where *n* = 200 and (*µ*_1_, *π*_1_) = (0, 0), (1, 0.1), (3, 0.1), (5, 0.1). Each black curve is from one sample realization. The red curve is the average of 50 black curves at each *γ* and the blue line is the underlying parameter value.

**Figure S2:**
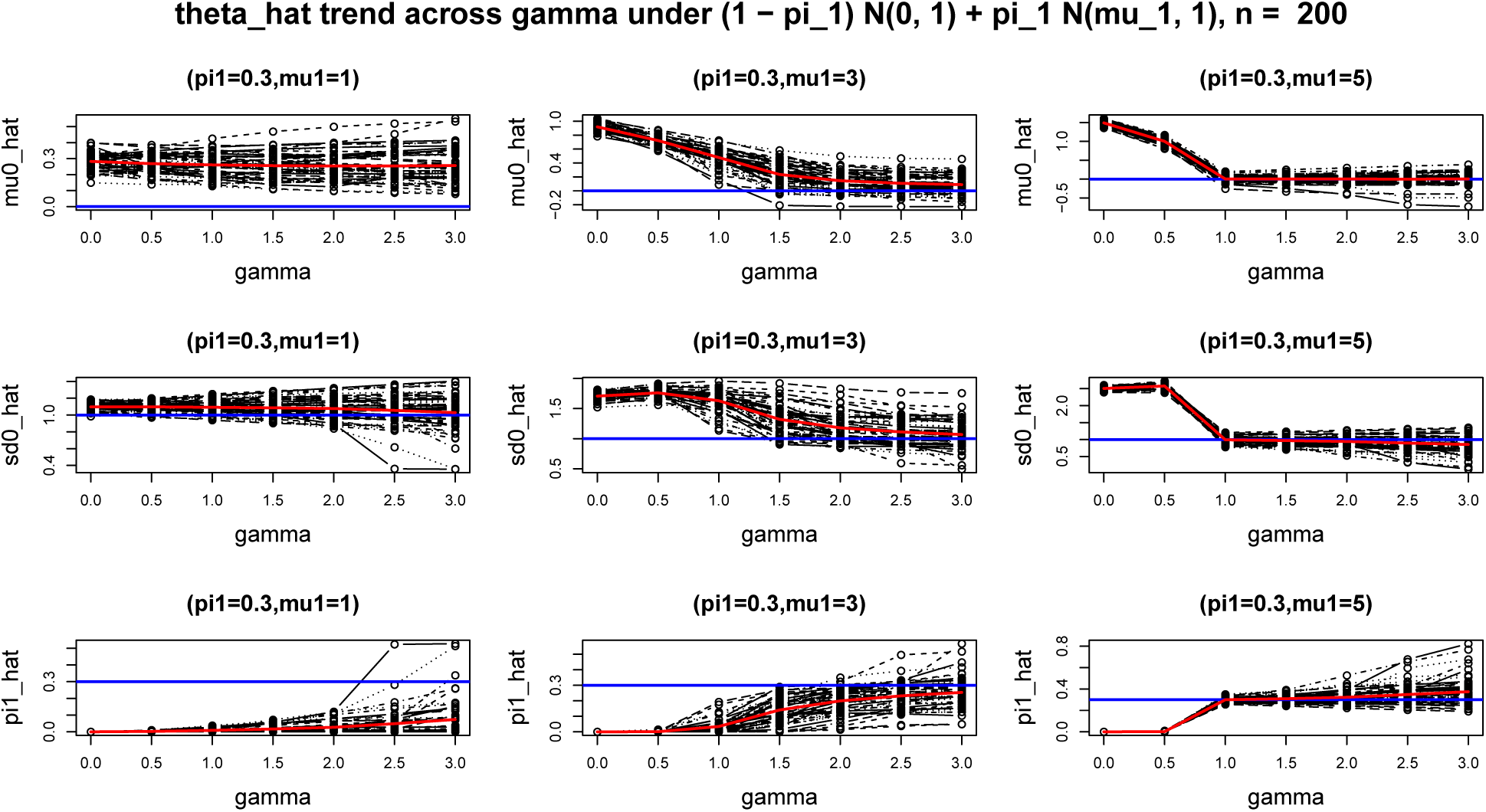
The same caption as Figure S1 under *n* = 200, (*µ*_1_, *π*_1_) = (1, 0.3), (3, 0.3), (5, 0.3).

**Figure S3:**
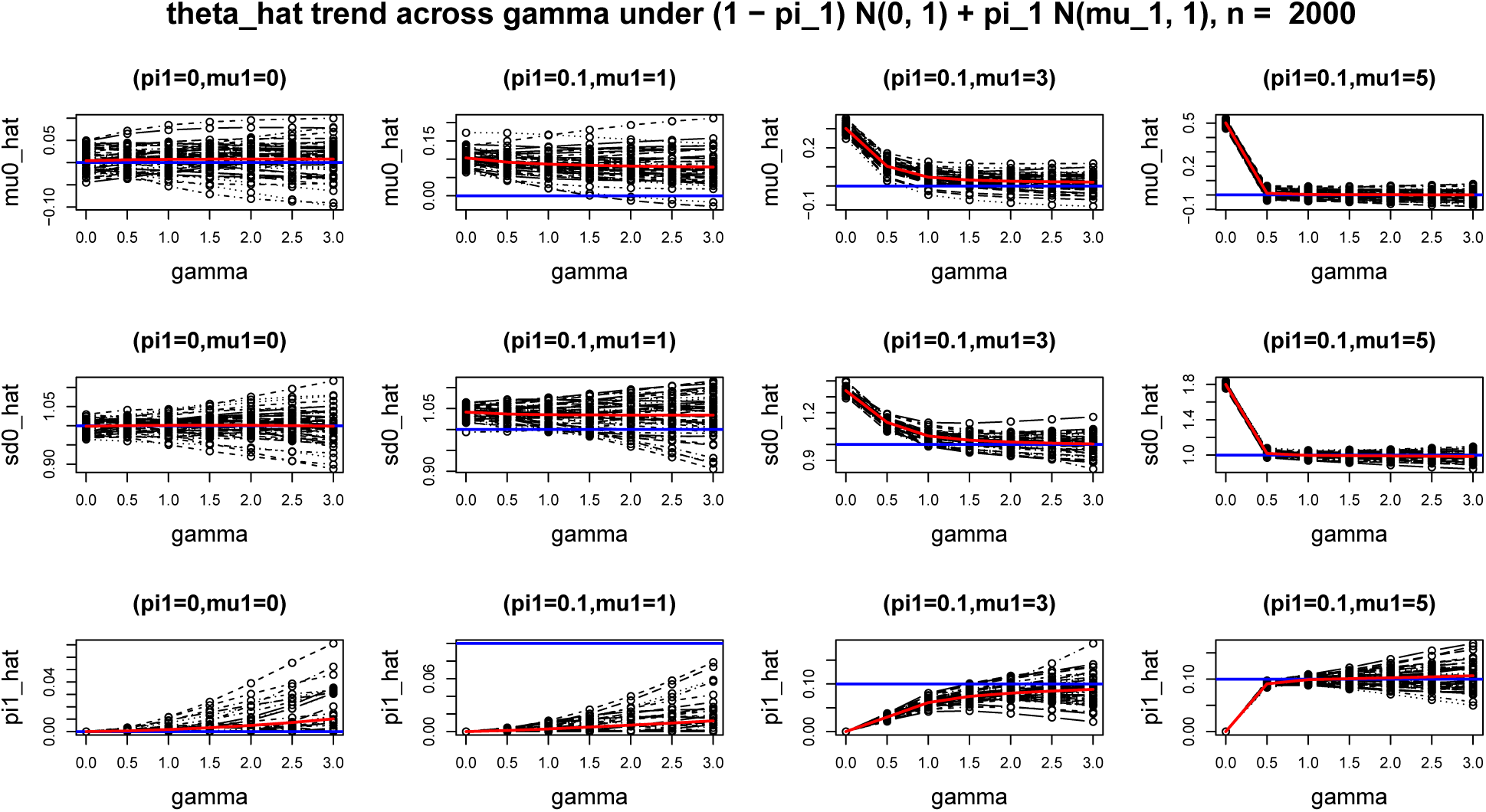
The same caption as Figure S1 under *n* = 2000, (*µ*_1_, *π*_1_) = (0, 0), (1, 0.1), (3, 0.1), (5, 0.1).

**Figure S4:**
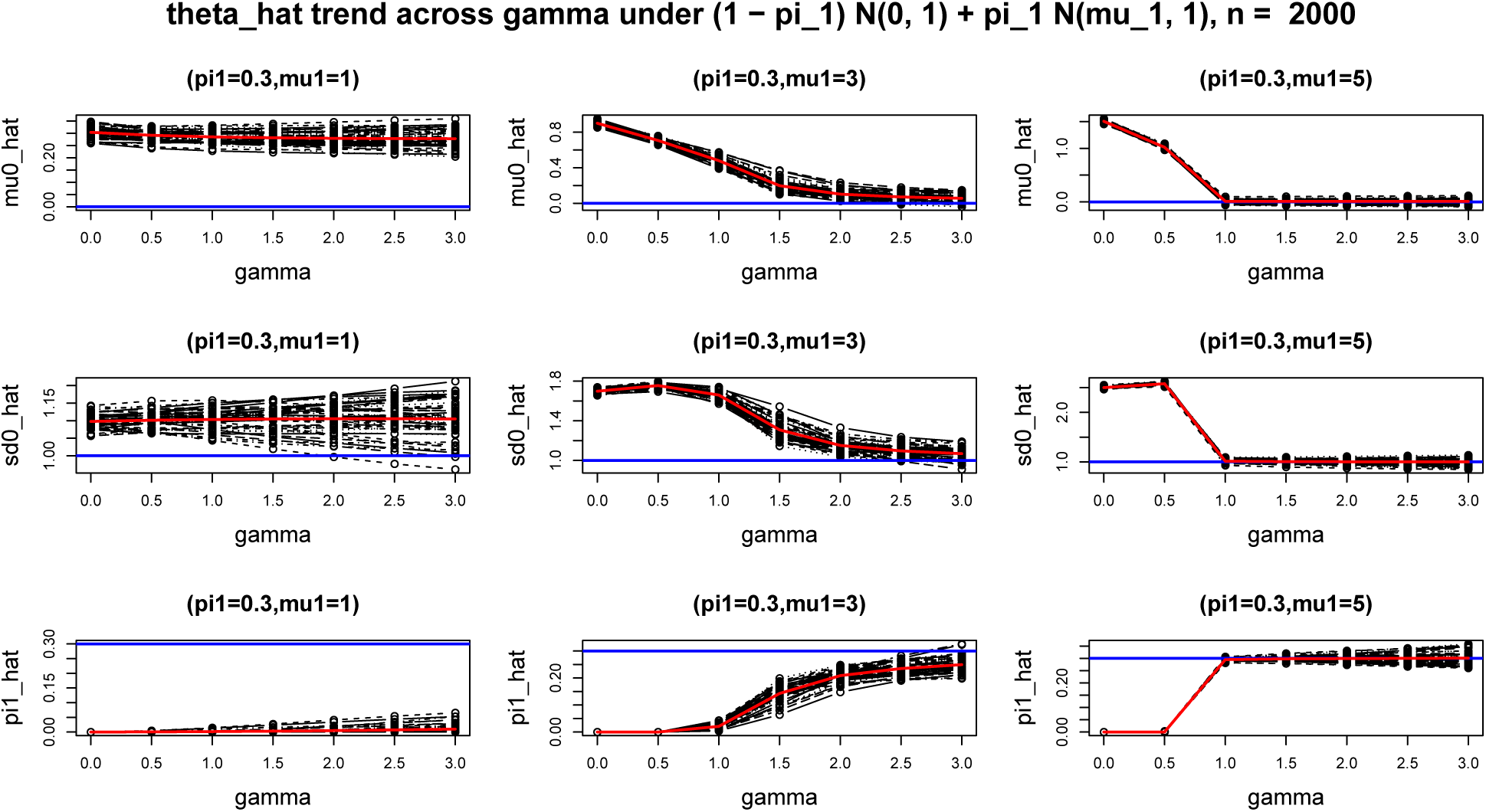
The same caption as Figure S1 under *n* = 2000, (*µ*_1_, *π*_1_) = (1, 0.3), (3, 0.3), (5, 0.3).

**Figure S5:**
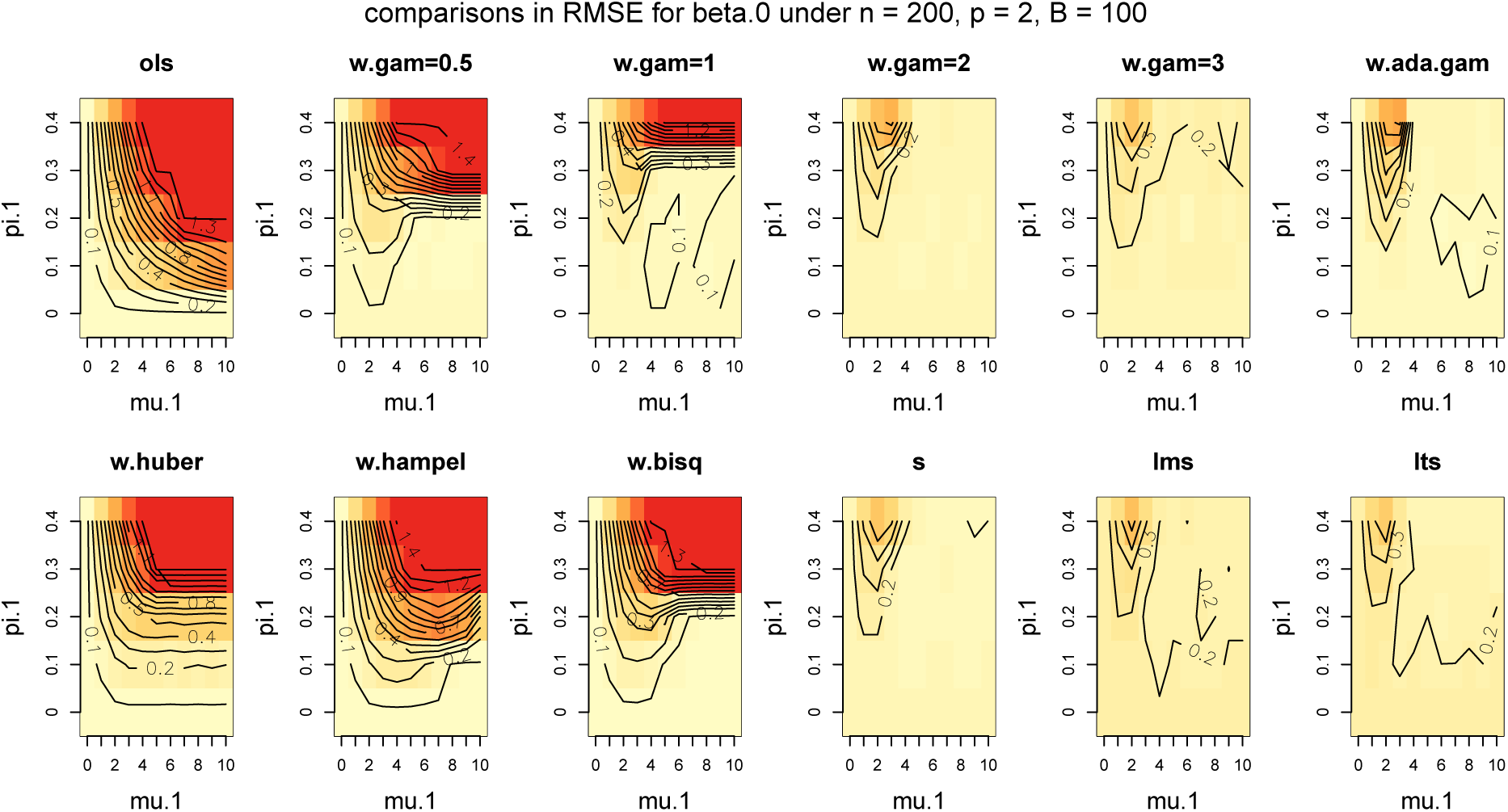
Heat map of RMSE comparisons for estimating ***β***_0_ under the regression model **y** = **X*****β***_**0**_ + ***ϵ*** where sample size *n* = 200 and *p* = 2, ***β***_0_ = **1**_**p×1**_ and 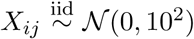 and the noise is i.i.d. from 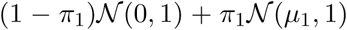, and (*π*_1_, *µ*_1_) = (0, 0) ∪ (0.1, 0.2, …, 0.4) × (1, 2, …, 10). The RMSEs are truncated by 2.

**Figure S6:**
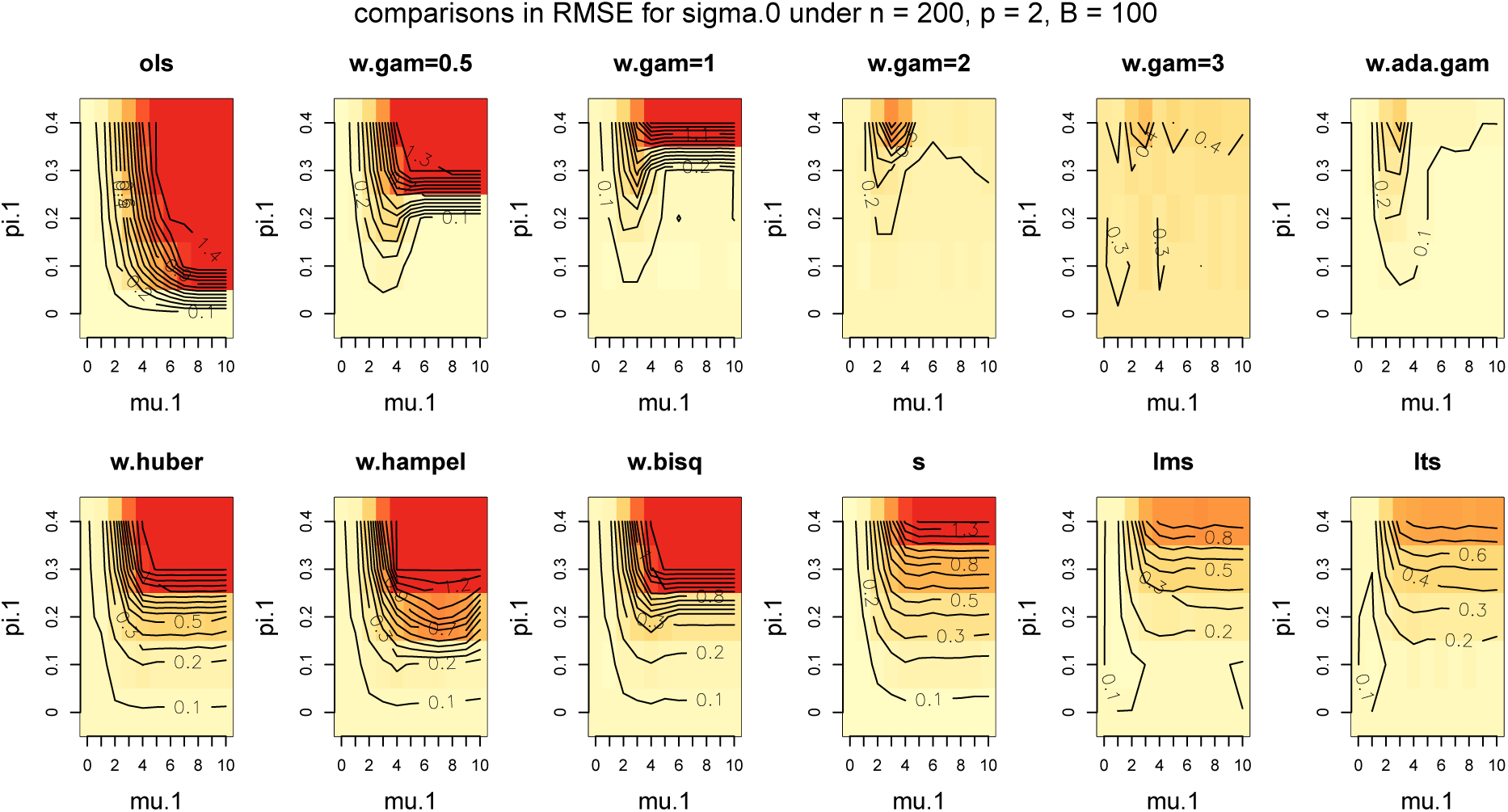
Heat map of RMSE comparisons for estimating *σ*_0_ under the same setting as in Figure S5.

**Figure S7:**
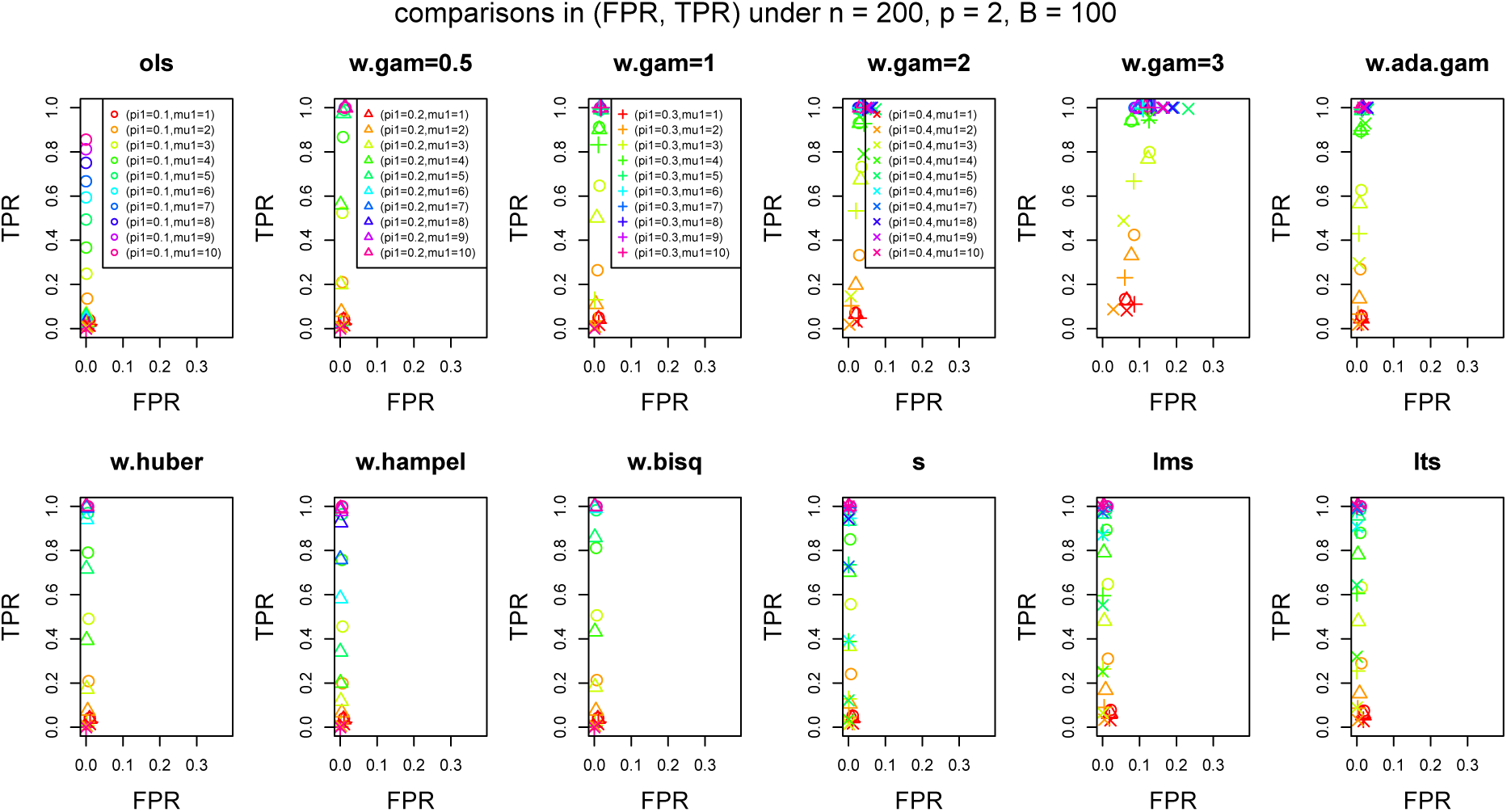
Scatter plot of (FPR, TPR) under the same setting as in Figure S5.

**Figure S8:**
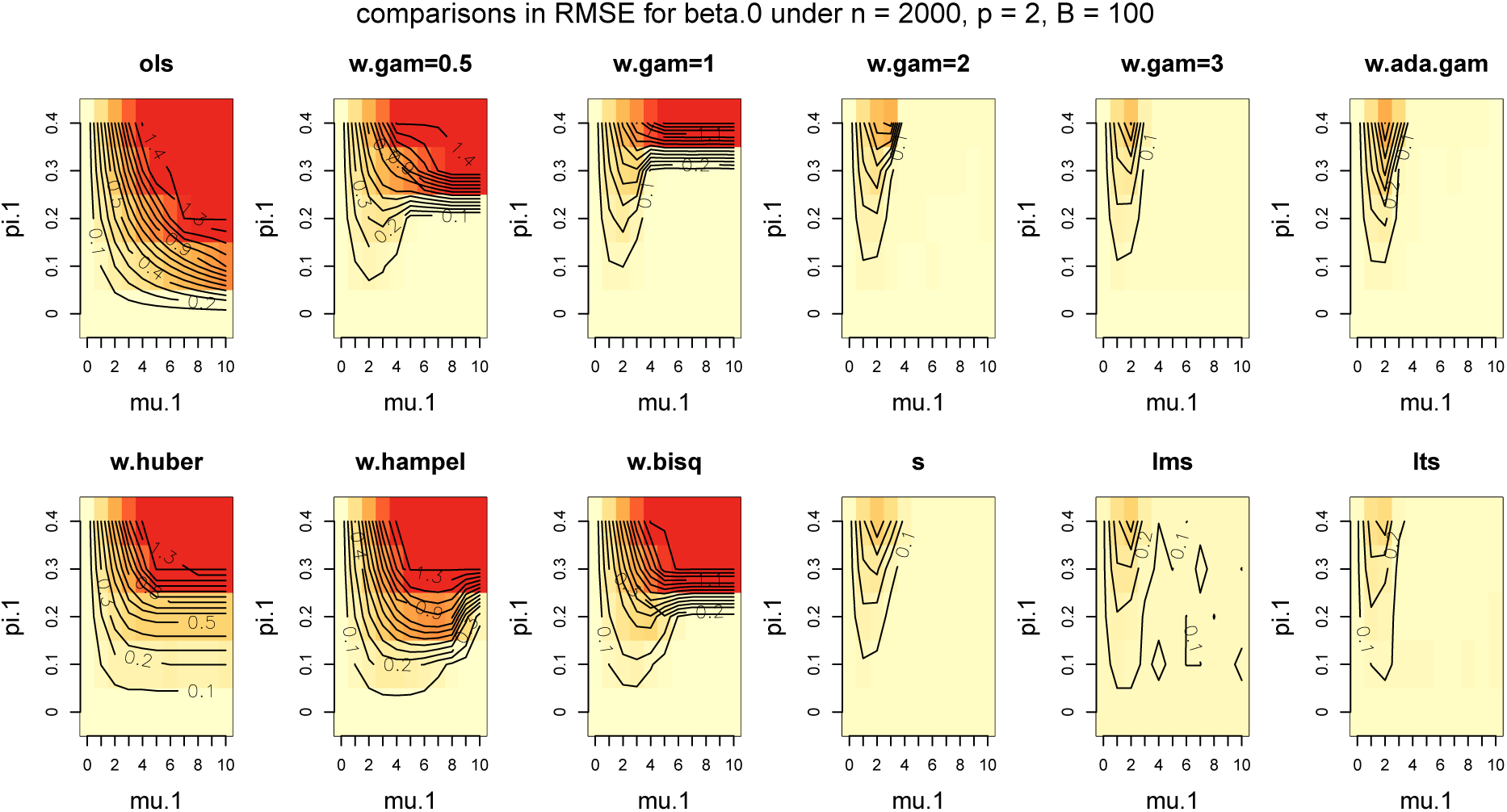
Heat map of RMSE comparisons for estimating ***β***_0_ in the same labels as in Figure S5 except *n* = 2000, *p* = 2.

**Figure S9:**
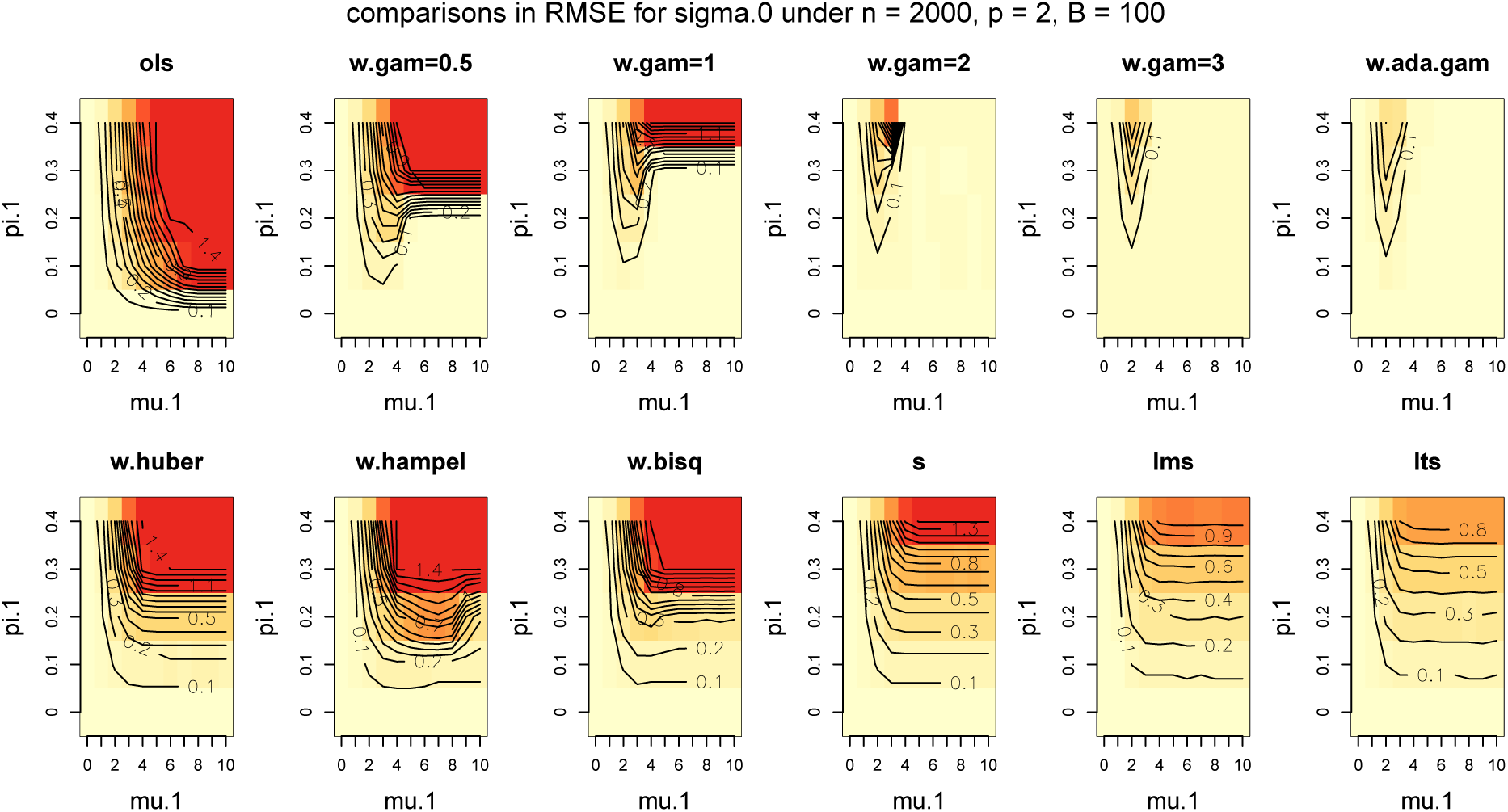
Heat map of RMSE comparisons for estimating *σ*_0_ in the same labels as in Figure S6 except *n* = 2000, *p* = 2.

**Figure S10:**
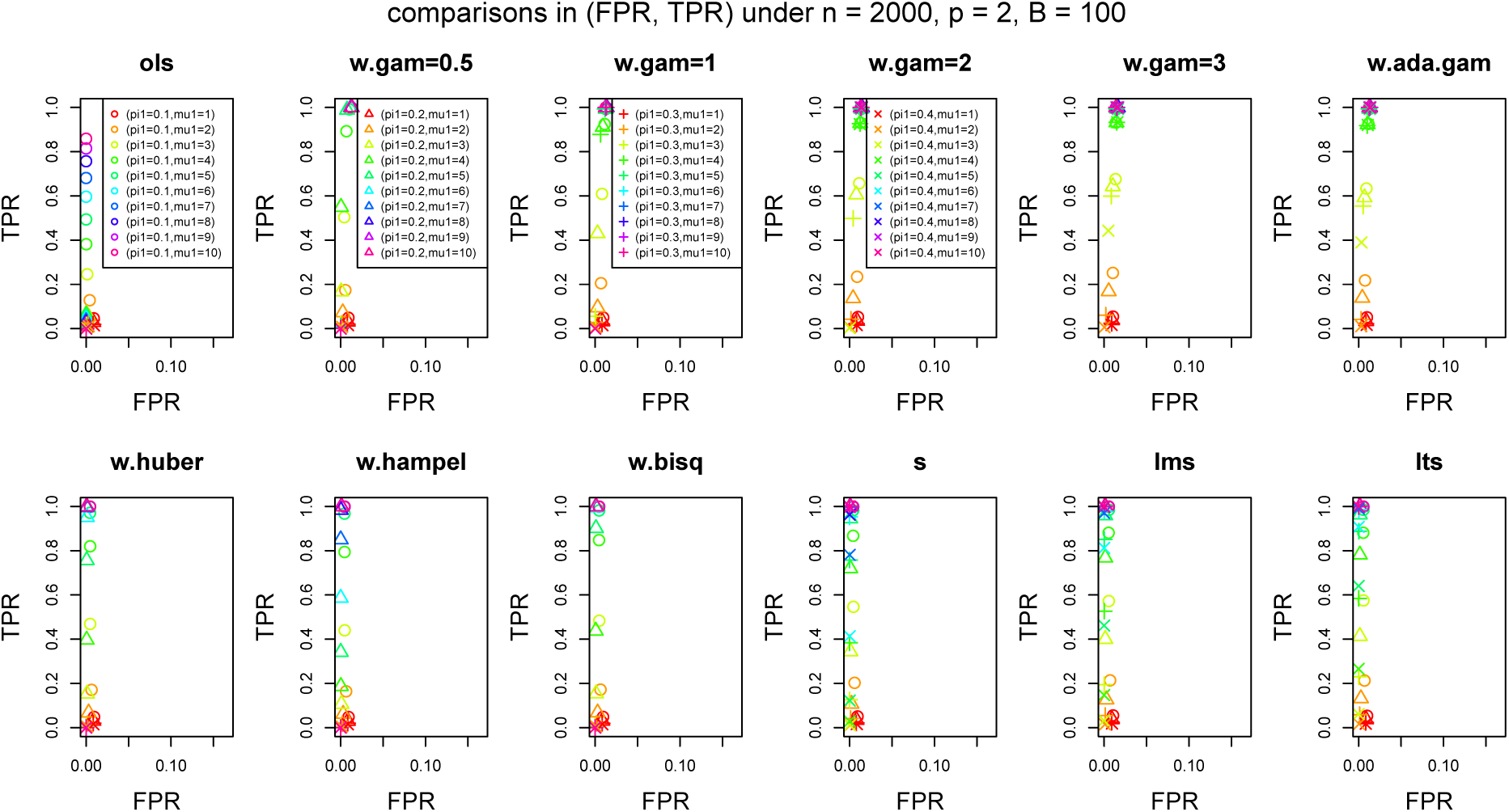
Scatter plot of (FPR, TPR) in the same labels as in Figure S7 except *n* = 2000, *p* = 2.

**Figure S11:**
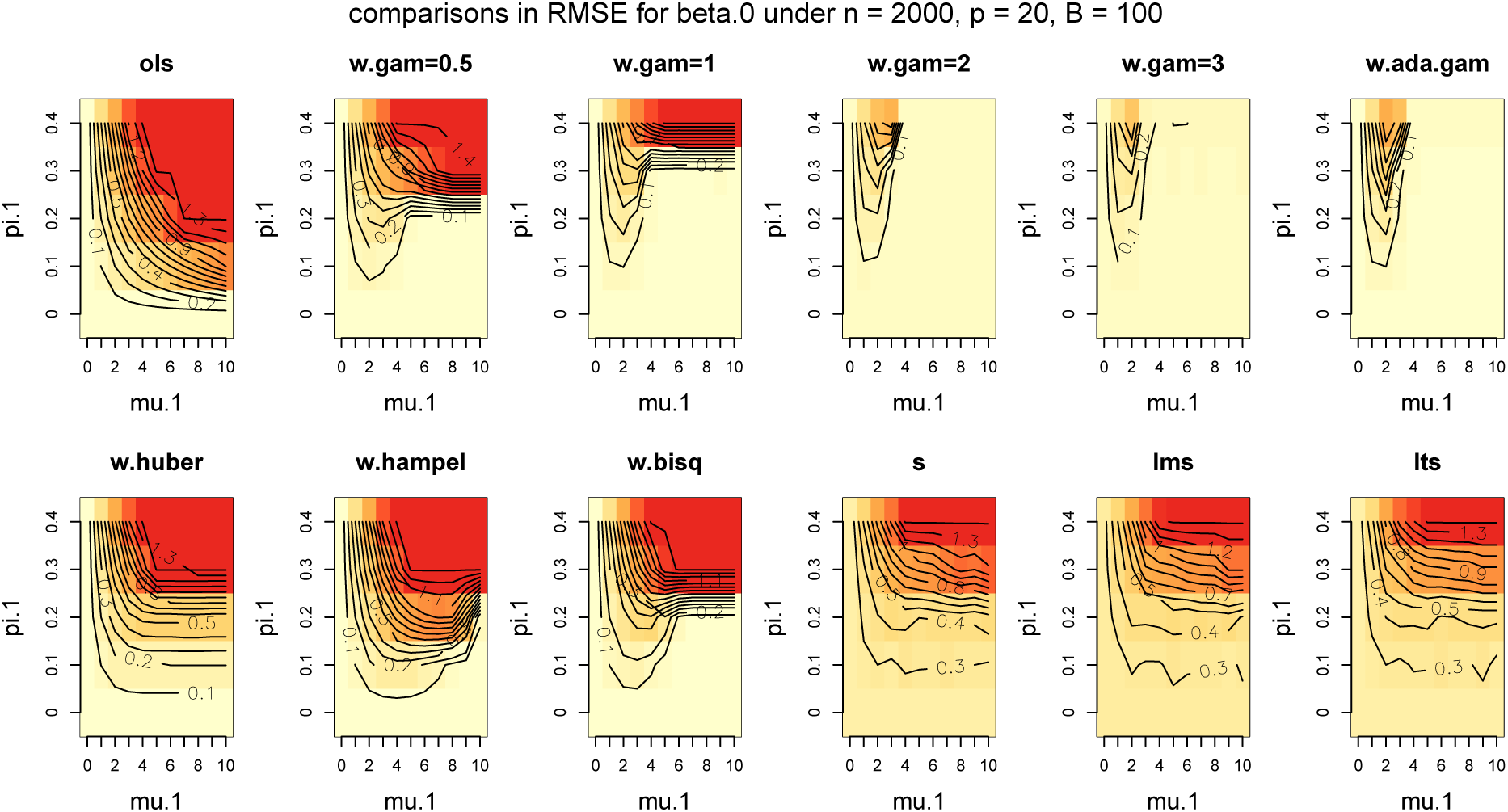
Heat map of RMSE comparisons for estimating ***β***_0_ in the same labels as in Figure S5 except *n* = 2000, *p* = 20.

**Figure S12:**
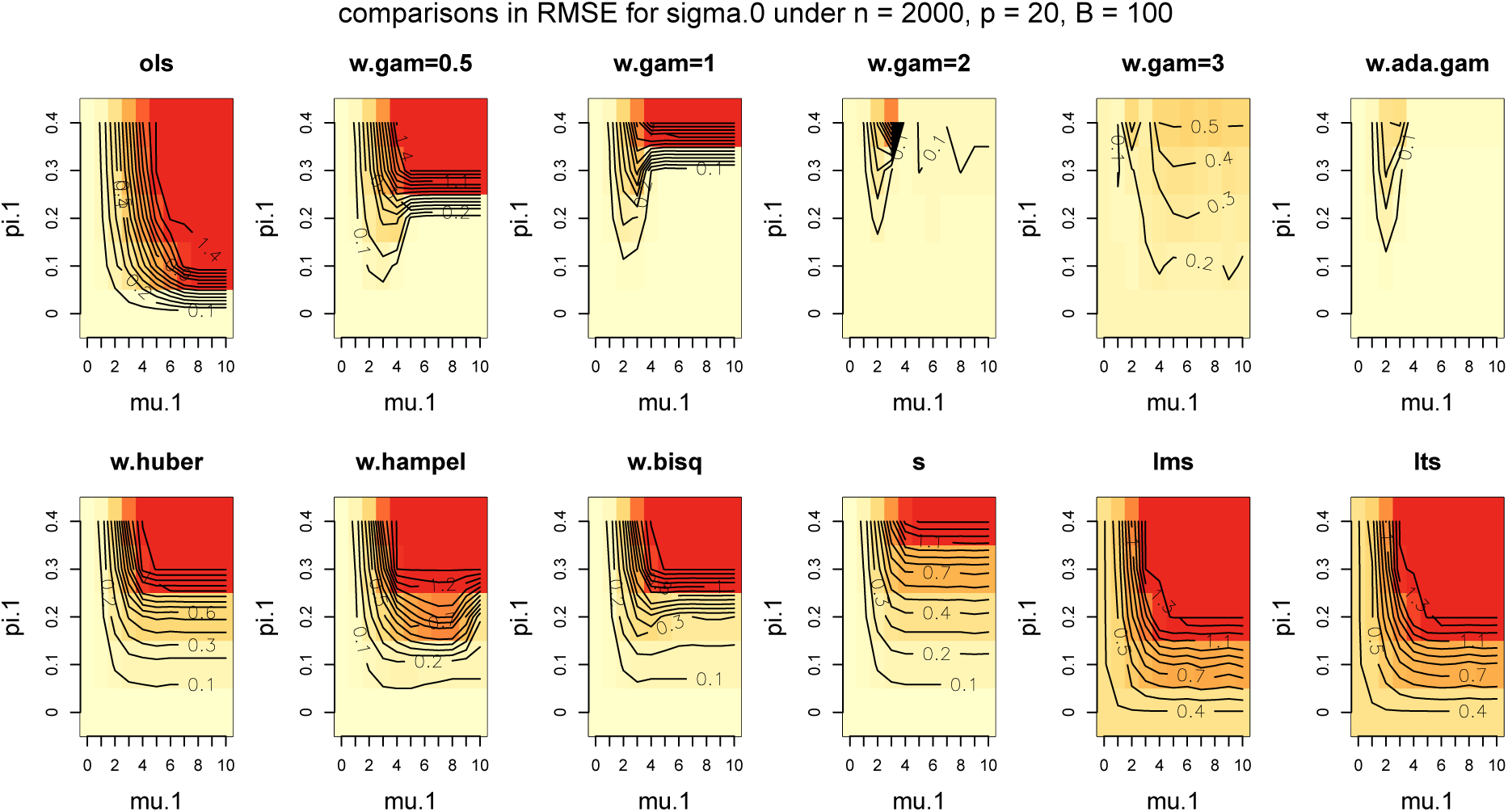
Heat map of RMSE comparisons for estimating *σ*_0_ in the same labels as in Figure S6 except *n* = 2000, *p* = 20.

**Figure S13:**
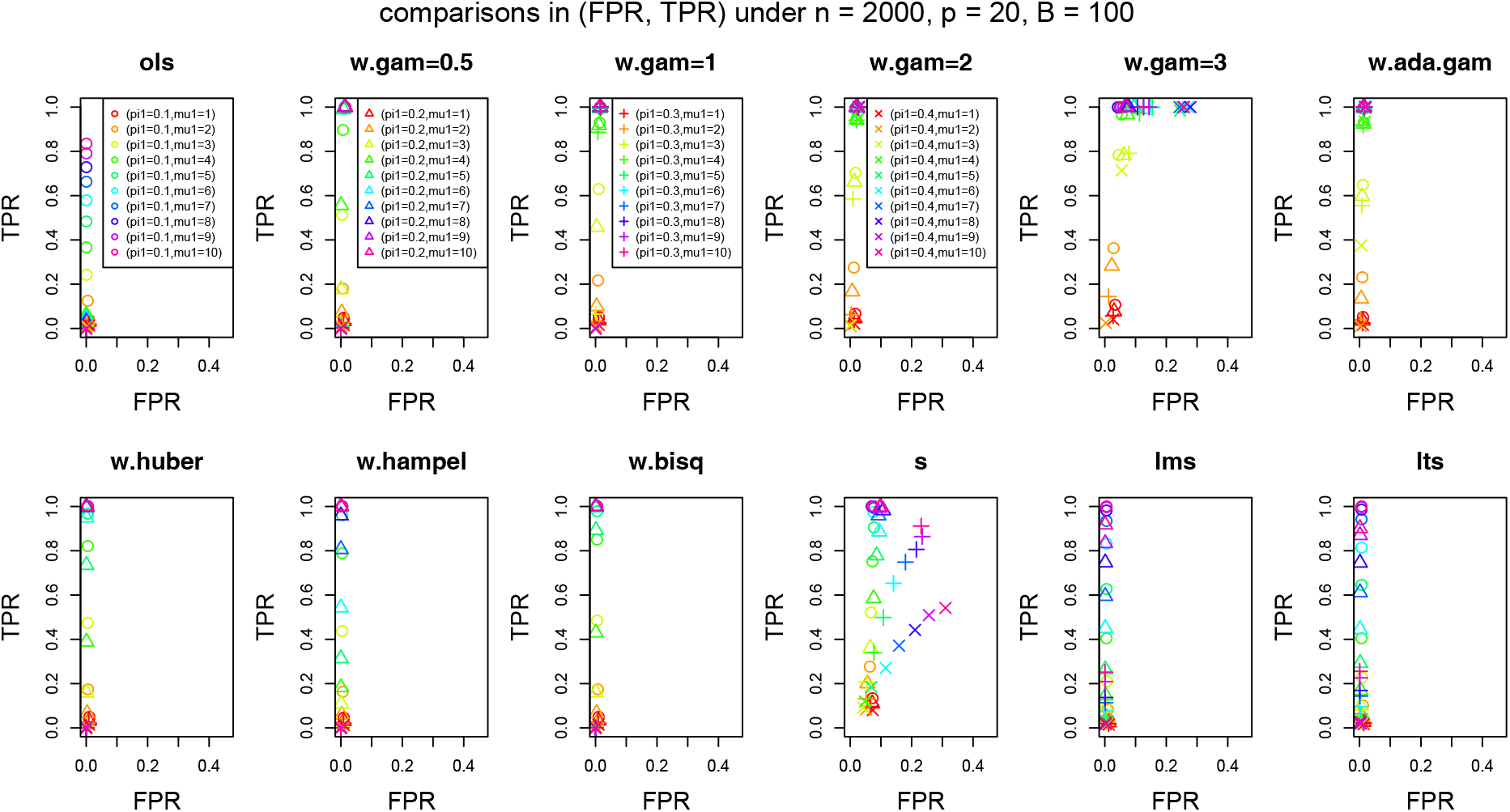
Scatter plot of (FPR, TPR) in the same labels as in Figure S7 except *n* = 2000, *p* = 20.

**Figure S14:**
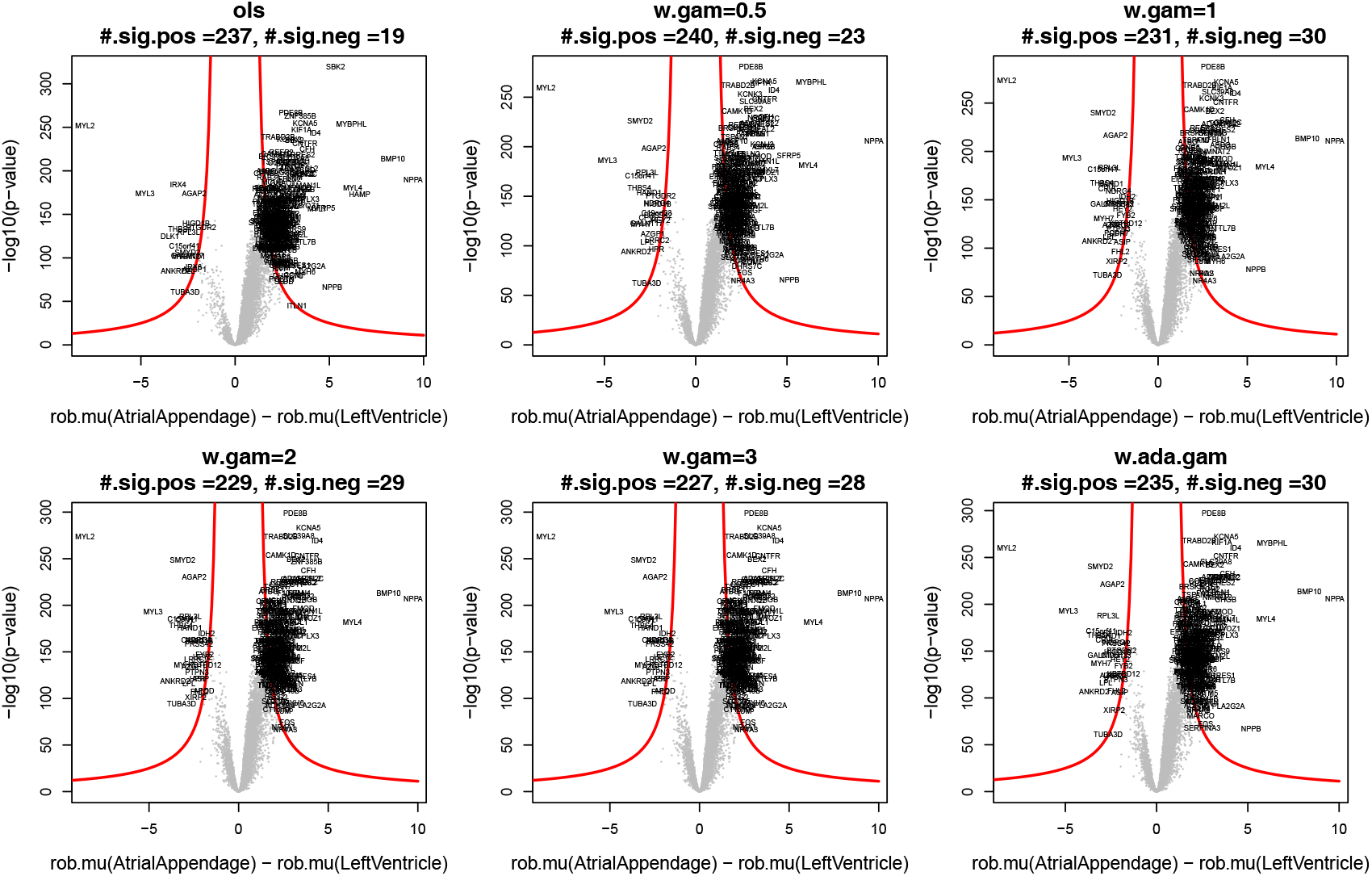
Volcano plot of minus of log_10_(*p*-value) versus log(fold change) 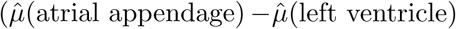) from two-sample *t*-test with unequal variances after removing the outliers from various *γ*-robustifying procedures. The red curve is hyperbolic cut with curvature parameter 100 and minimum fold change parameter 1 (Singh et al., 2016). The gene names are labeled if they are above the red curve.

